# Targeting CD73-A_2a_R-Mediated Adenosine Signaling at the Tumor-Immune Interface Overcomes Radioresistance

**DOI:** 10.64898/2026.05.26.727904

**Authors:** Shruti Bansal, Luis Aparicio, Aparna Krishnan, Chang Liu, Lindsay Caprio, Anna M Chiarella, Samanta Sarti, Jason Piersant, Chelsea L. Rahiman, Julia An, Patrick Mccann, Namita Sen, Brian E. Ragaishis, Fatemeh Derakhshan, Bret Taback, Anil Rustgi, Benjamin Izar, Catherine S. Spina

## Abstract

**Background:** Radiotherapy efficacy is constrained by an immunosuppressive tumor microenvironment (TME) enriched in extracellular adenosine and suppressive myeloid populations that attenuate cytotoxic T-cell responses. The CD73–adenosine–A_2a_/A_2b_ receptor axis represents a key metabolic immune checkpoint; however, the relative contributions of tumor cell–intrinsic versus host-derived adenosine signaling to radiotherapy response remain incompletely defined.

**Methods:** Using orthotopic murine breast carcinoma models, we interrogated radiation-induced adenosine dynamics and downstream immune remodeling through quantitative adenosine measurements, bulk RNA sequencing, and multiparameter flow cytometry. Genetically engineered models were employed to dissect the roles of tumor-derived CD73 and host A_2a_/A_2b_ receptors in regulating radiosensitivity. Therapeutic studies evaluated combinatorial targeting of CD73 and A_2a_/A_2b_ receptors with radiotherapy and anti–PD-1, followed by comprehensive immune profiling in breast carcinomas.

**Results:** Tumor cell–intrinsic CD73 and host A2A receptor signaling cooperatively drive radioresistance and tumor progression. Radiotherapy induces a rapid surge in intratumoral adenosine, triggering transcriptional and cellular programs consistent with myeloid-mediated immunosuppression and lymphocyte dysfunction. Although T-cell infiltration increases at later time point post-irradiation, effector function remains constrained. Pharmacologic inhibition of CD73 and A_2a_/A_2b_ receptors partially restores T-cell functionality but is insufficient for durable tumor control as monotherapy. In contrast, concurrent blockade of adenosine signaling during radiotherapy, followed by adjuvant PD-1 inhibition, amplifies adaptive antitumor immunity and significantly enhances tumor control.

**Conclusions:** These findings define a mechanistic link between radiation-induced adenosine signaling and immune dysfunction in the TME. Targeting the CD73–A_2a_/A_2b_ axis in combination with radiotherapy and checkpoint blockade represents a rational strategy to overcome radioresistance and improve antitumor immunity.

**STATEMENT OF SIGNIFICANCE:** The tumor and immune cell contributions to adenosine signaling play a central role in shaping the therapeutic outcomes of tumor irradiation. Therapeutic targeting of the adenosine signaling axis improves radiosensitivity and efficacy of checkpoint blockade.

## INTRODUCTION

Radiotherapy is offered to approximately half of cancer patients. Depending on the dose and schedule of delivery, tumor irradiation can promote adaptive anti-tumor immunity (1, 2) and produce immunosuppressive cues in the tumor microenvironment (TME) (3). Radiation causes the release of adenosine triphosphate (ATP) from damaged or dying cells in the TME (4). ATP is quickly metabolized to adenosine through a series of ectonucleotidases including hydrolysis by CD39 (*ENTPD1*) to ADP and AMP and subsequently from AMP to adenosine by CD73 (*NT5E*). Meanwhile, radiation-induced DNA damage results in production of cyclic GAMP (cGAMP) that is hydrolyzed to AMP by ENPP1 and to adenosine by CD73 (5). Adenosine mediates its immunosuppressive functions through four adenosine receptors with Adora 2a receptor (A_2A_R) and Adora 2b receptor (A_2B_R) being the two predominant members (5).

Adenosine signaling suppresses a diverse range of immune cell functions. A_2A_R signaling in T cells exerts its suppressive effect by decreasing T cell receptor (TCR) signaling, CD28 co-stimulation and IL-2 receptor signaling, thereby blocking T cell activation, proliferation and secretion of cytokines IFNγ, IL-2, IL-6, and TNFα (5,6). A_2A_R signaling in CD4+ T cells promotes their differentiation into Tregs and increases the secretion of immunosuppressive cytokines (7). In neutrophils, A_2A_R signaling inhibits oxidative burst, matrix metalloproteinase secretion and trans-endothelial migration (8). A_2A_R and A_2B_R signaling suppress antigen presentation by dendritic cells (DCs) and macrophages by downregulating MHC II expression, co-stimulation by decreased expression of CD86 and secretion of pro-inflammatory cytokines IL-12 and TNFα while promoting production of pro-tumor factors IL-6, IL-10, TGFβ and VEGF (9).

There is mounting evidence that supports the prognostic significance and therapeutic value of inhibiting adenosine signaling in cancer. Expression of CD73 has been shown to be prognostic across multiple cancer types (10, 11, 12). In the preclinical model of breast cancer, the combination of CD73 inhibition and tumor irradiation enhances antigen presentation and improves tumor control. When combined with an immune checkpoint inhibitor, anti-CTLA-4, CD73 blockade reduced metastatic disease burden, and improved overall survival (13). In the clinical setting, phase I/II SYNERGY trial, the addition of anti-CD73 (oleclumab) to chemoimmunotherapy for patients with metastatic TNBC failed to show benefit (14). However, phase 2 COAST trial demonstrated that for patients with unresectable stage III non-small cell lung cancer (NSCLC), addition of CD73 blockade to adjuvant anti-PD-L1 after definitive chemoradiation increased the overall response rate (ORR) and prolonged progression-free survival (PFS), demonstrating clinical value of CD73 blockade after tumor irradiation (15). A similar benefit was observed in the phase II Neo-CheckRay study combining CD73 inhibition to neoadjuvant chemotherapy (NACT), stereotactic body radiotherapy (SBRT) with and without adjuvant anti-PD-L1 for patients with luminal B breast cancer. They demonstrated that 35.6% of patients who received anti-PD-L1 and CD73 blockade with chemoradiation achieved a pathologic complete response (pCR) compared with 17.8% patients who received chemoradiation alone (16).

Others have targeted A_2A_R to mitigate downstream signaling of adenosine to alleviate immune suppression. In a murine model of TNBC, co-administration of immunogenic cell death inducer and A_2a_R antagonist promoted maturation of dendritic cells (DCs) and enhanced CD8^+^ T cells infiltration while reducing granulocytic myeloid-derived suppressor cells (G-MDSCs) infiltration thus improving antitumor immunity (17). To date, there has been limited success with A_2A_R inhibition in the monotherapy setting with modest activity when combined with anti-PD-L1 (18). Hence, other strategies are needed to sensitize tumors to adenosine signaling modulation for improved responses.

Radiotherapy induces immunogenic cell death and tumor control while activation of CD73-A_2a_R signaling triggers immunosuppression impairing anti-tumor immunity after radiation. The extent to which CD73–A_2a_R signaling represents a targetable mechanism of adaptive radioresistance remains incompletely defined. Herein, we dissect the contribution of adenosine signaling proteins-CD73 and A_2a_R expressed by tumor and immune cell compartments towards radioresistance and test the clinical value of inhibiting adenosine signaling pathway to improve radiosensitivity and efficacy of checkpoint blockade.

## METHODS

### Mice

Female C57BL/6 (RRID: IMSR_JAX:000664), BALB/c (RRID: IMSR_JAX:000651), CD73^-/-^, LysM^Cre^ (RRID: IMSR_JAX:004781), A_2a_R^fl/fl^ (RRID: IMSR_JAX:010687) mice were purchased from the Jackson laboratory (Bar Harbor, Maine, USA). A_2a_R^-/-^A_2b_R^-/-^ mice were a kind gift from the laboratory of Dr. Stephen Tilley at University of North Carolina at Chapel Hill. LysM^Cre^A_2a_R^fl/fl^ were generated by crossing A_2a_R^fl/fl^ mice with LysM^Cre^ mice. Mice were used between 8 and 10 weeks of age. All animals were housed as per the guidelines of American Association of Laboratory Animal Care regulations. All experiments were carried out after approval by the Animal Ethics Committee of Columbia University. Based on a power analysis conducted to achieve 80% probability of detecting a true effect we used 10-12 mice in each experiment.

### Cell lines

Murine luminal B breast cancer cell line EO771 (ERα-, ERβ+, PR+, HER2+) and triple negative breast cancer cell line 4T1 (ER-, PR-, HER2-) was procured from ATCC. Knockdowns were generated in EO771 cells by depleting CD73, Enpp1 or both using the CRISPR-Cas9 system. The detailed media for maintenance of these cell lines is outlined in supplemental methods.

### Patient samples

Breast tumor and adjacent normal breast tissue were obtained from untreated patients undergoing biopsy or surgery at CUIMC under an Institutional Review Board (IRB) approved protocol (supplementary table 1). Half of the tissue was collected in tissue storage solution (TSS, Miltenyi Biotech) supplemented with adenosine inhibitors-40nM dipyridamole (DPP, Sigma Aldrich) and 100nM erythro-9-(2-hydroxy-3-nonyl) adenine (EHNA, Sigma Aldrich) and used for adenosine quantification. The second half of the tissue was fixed for immunohistochemistry (IHC). The detailed procedure for IHC is included in supplemental methods.

### Adenosine quantification

Adenosine was measured by high-performance liquid chromatography-tandem mass spectrometry (UPLC-MS/MS) on a platform comprising of Waters Xevo TQA triple quadrupole mass spectrometer integrated with a Waters Aquity Premier UHPLC (Waters, Milford, MA). Adenosine was extracted from plasma and tissue homogenates spiked with inhibitors and deuterated internal standard as described in Sharma et al (30) with modifications. Chromatographic separation was done on a Waters Aquity UPLC CSH Phenyl Hexyl column (1.7µm, 2.1X100mm) maintained at 50°C employing gradient elution using water and acetonitrile with 0.1% formic acid as mobile phase. Positive ESI-MS/MS mass spectrometry under MRM mode was performed using the following transitions: adenosine 268.2>136.1 and 13C5 adenosine 273.2>136.1.

### Tumor model and treatment injections

EO771 or 4T1 cells (ATCC Cat# CRL-2539, RRID:CVCL_0125), 5x 10^5^ cells/100ul were implanted orthotopically in the mammary fat pad of mice. Tumor volume measurements were calculated by multiplying length x width x height. Mice were then randomized to different groups such that average tumor volume were similar in each group. Mice tumors were treated with radiation (8Gy x 1 or 8Gy x 3 daily) using the X-RAD 320 (self-contained X-ray system designed for precise radiation dosage delivery to small animals). Mice were administered AB680 (quemliclustat, CD73 inhibitor, Arcus Biosciences) via oral gavage, AB928 (etrumadenant, A_2a_R/A_2b_R antagonist, Arcus Biosciences) via subcutaneous injection and AB122 (aPD-1, zimberlimab, Arcus Biosciences) or rat IgG2a isotype antibody (Arcus Biosciences) via intra-peritoneal injection as mentioned in figures.

### Statistics

Significant differences between experimental groups were determined using a two-tailed Student *t*-test (to compare two samples), an ANOVA analysis followed by Tukey’s multiple comparisons test (to compare multiple samples) or Mann Whitney test (non-parametric test) in GraphPad Prism 6 (La Jolla, CA, RRID:SCR_002798). For all analyses, a *P* value <0.05 was significant.

## RESULTS

### Expression of CD73 on cancer cells and A_2a_R on immune cells drives adenosine signaling in TME

Adenosine is known to be higher in tumors compared to normal tissues (19), thereby contributing to the suppressive milieu of the TME. We observed a 3-fold increase in intratumoral adenosine in primary breast tumors from humans and mice compared to paired normal breast tissue (Figure 1A, 1B). To understand the distribution of CD73 and A_2a_R in the TME, we performed immunohistochemistry (IHC) of human breast cancer tissue. We demonstrated that CD73 is localized to tumor cell membranes (H- score of 100) where 90% stained weakly (1+) and 5% stained strongly (2+) (Figure 1C, 1D). On the other hand, A_2a_R was more highly expressed by immune cells (H-score of 270) where 90% stained strongly (3+) (Figure 1C, 1E). High expression of CD73 by breast cancer cells and A_2a_R by immune cells (Figure 1D, 1E) suggests a potential cooperative relationship between the two cellular compartments. Quantification of transcriptional activity of the adenosine signaling pathway genes (*Enpp1*, Entpd1 (*CD39)*, Nt5e (*CD73)*, *adora2a* (A_2a_R) and *adora2b* (A_2b_R) in murine luminal B breast cancer cells, EO771, and triple negative breast cancer cells (TNBC), 4T1, *in vitro* confirmed that CD73 is more highly expressed than A_2a_R (Figure 1F). Immunophenotyping of orthotopic EO771 and 4T1 tumors revealed that breast TME was heavily enriched in CD11b+ myeloid cells making up 85-90% of the CD45+ immune compartment. T cells are significantly less abundant (5-7%) and B cells, NK cells, dendritic cells were less than 2% (Figure 1G, 1H). The myeloid rich breast TME of EO771 tumors is comprised of a mix of macrophages (31%), PMNs (22%) and monocytes (21%) (supplementary figure S1A) whereas in 4T1 tumors, the myeloid compartment is predominately PMNs (72%, supplementary figure S1B). We quantified the expression of the CD39, CD73 and Enpp1 and A_2a_R in the immune (CD45^+^) and non-immune (CD45^-^) compartments of EO771 and 4T1 tumors by flow cytometry (Figure 1I, 1J). We found that CD73 was the most abundant in the CD45^-^ compartment (18-20% of population) of EO771 and 4T1 tumors (Figure 1I, 1J). Whereas, in the CD45^+^ compartment of EO771 and 4T1 tumors, higher proportion of CD11b+ cells expressed CD39 (63.5%, 74%), A_2a_R (32.5%, 39%), CD73 (22.3%, 63%), enpp1 (12.75%, 29%) compared to T cells (CD39: 3.81%, 1.5%; A_2a_R: 0.16%, 0; CD73: 2.26%, 3%; enpp1: 0.43%,0) and minimal (<1%) in NK cells, B cell, and DCs (Figure 1K, 1I). However, in the CD11b+ compartment of EO771 tumors, adenosine signaling pathway proteins were most expressed by macrophages (CD39: 26.4%, A_2a_R: 20.8%, CD73: 10.8%, enpp1: 4.8%), with lower abundance in monocytes and PMNs (supplementary figure S1C). In 4T1 tumors, adenosine signaling pathway proteins were predominantly expressed by PMNs (CD39: 51%, A_2a_R: 19.5%, CD73: 42.5%, enpp1: 38.4%, supplementary figure S1D). In summary, CD73 is most abundant in the CD45^-^ compartment, underscoring the role of the non-immune compartment, including tumor, in the production of suppressive adenosine. Although heterogeneous in the composition of myeloid cells subsets, overall CD11b^+^ myeloid cells in both EO771 and 4T1 tumors had the most abundant expression of A_2a_R, compared to other cellular compartments, suggesting a preferential role in promoting the downstream suppressive adenosine signaling.

**Figure 1:**
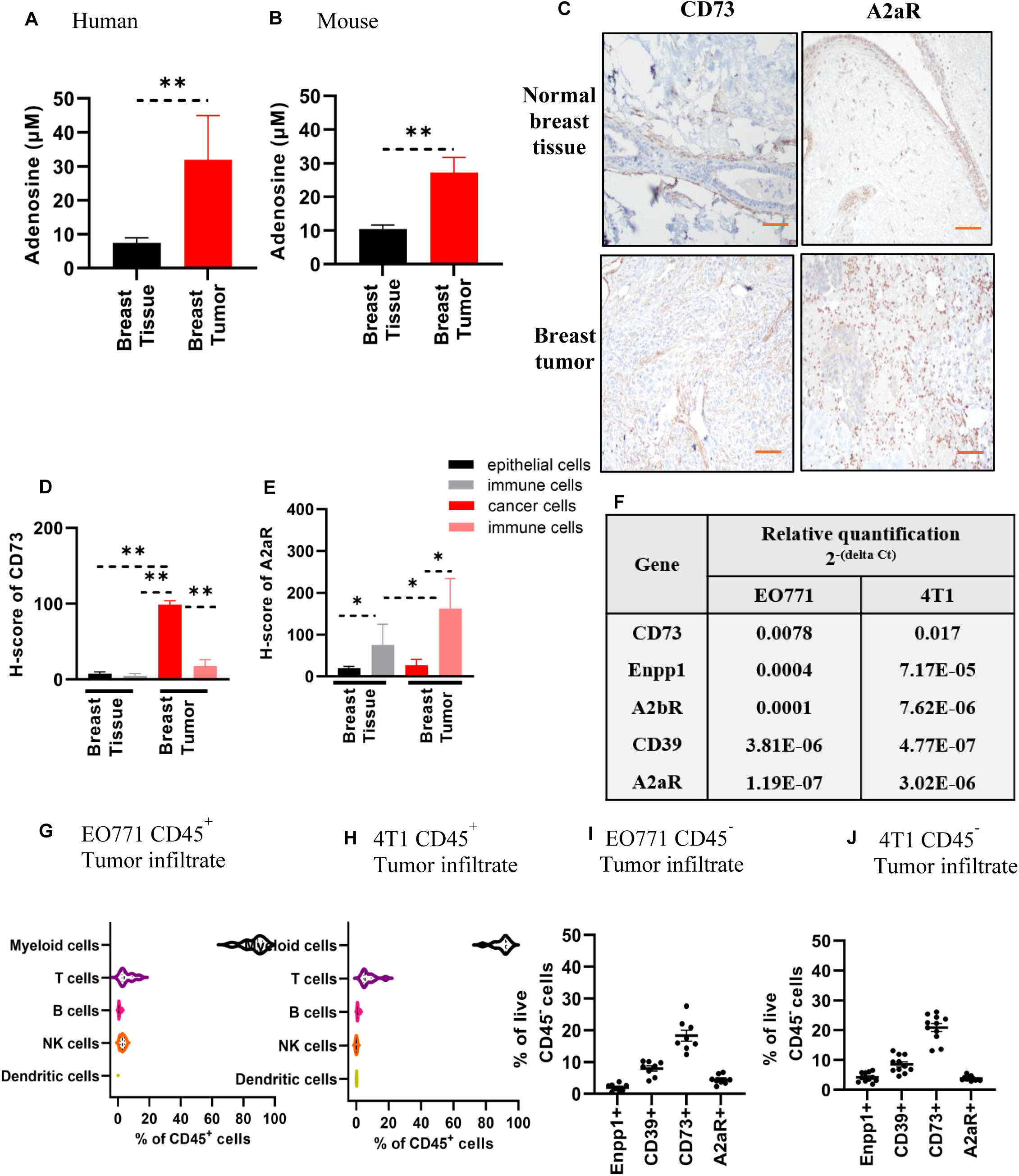
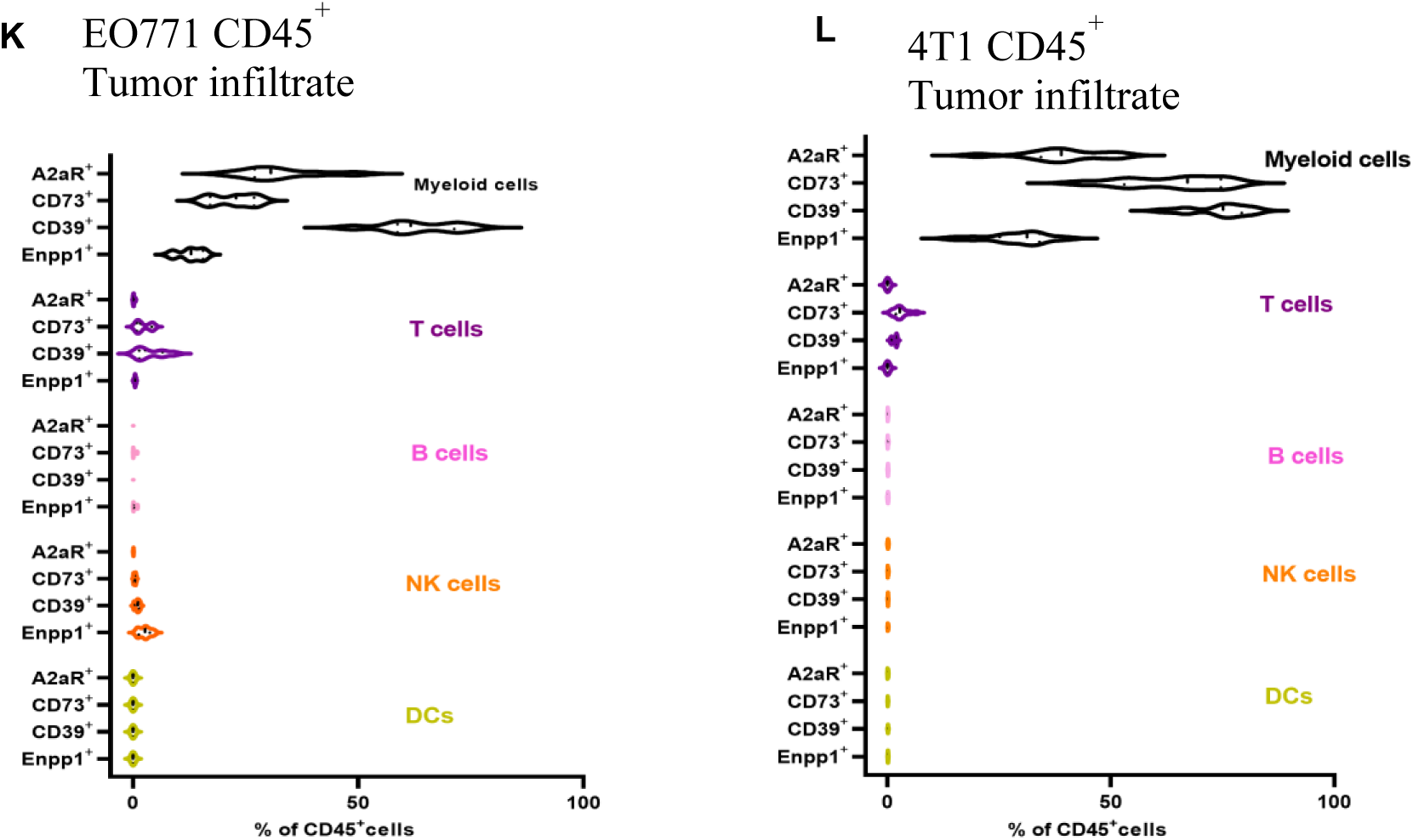
Intra-tumoral adenosine concentration (Mean ± SEM) in (A) human and (B) murine breast tumors and normal breast tissue quantified by LC-MS. (C) IHC staining: Magnification 200 X, Scale 100 μM and (D, E) H-score (Mean ± SEM) of CD73 and A_2a_R in human normal breast tissue (epithelial cells, immune cells) and breast tumors (cancer cells, immune cells). (F) Relative expression of genes in the adenosine signaling axis in EO771 and 4T1 murine breast cancer cells compared to GAPDH by qPCR. (G) Frequency of immune cell subtypes (Mean ± SEM) in EO771 tumors by spectral flow cytometry. (H) Frequency of immune cell subtypes (Mean ± SEM) in 4T1 tumors by spectral flow cytometry. (I) Abundance of adenosine signaling proteins (Mean ± SEM). in CD45^-^ compartment of EO771 tumors. (J) Abundance of adenosine signaling proteins (Mean ± SEM). in CD45^-^ compartment of 4T1 tumors. (K) Expression of adenosine signaling proteins on CD45^+^ immune cells (Mean ± SEM) in EO771 tumors quantified using spectral flow cytometry. (L) Expression of adenosine signaling proteins on CD45^+^ immune cells (Mean ± SEM) in 4T1 tumors quantified using spectral flow cytometry. Tumors were harvested from n=10-12 mice, 25 days post-implantation. (myeloid cells (CD45^+^CD11b^+^TCRβ^-^), T cells (CD45^+^CD11b^-^TCRβ^+^), B-cells (CD45^+^CD11b^-^CD19^+^), dendritic cells (CD45^+^CD11b^-^CD11c^+^CD103^+^), NK cell – natural killer cells (CD45^+^CD11b^-^CD11c^-^NK1.1^+^).

### Radiotherapy induces immunosuppressive cues in TME acutely post-treatment

We hypothesized that radiation generates a metabolic checkpoint- adenosine and alters myeloid and lymphoid cell populations in the TME thus dampening anti-tumor immunity. Indeed, in our EO771 breast cancer model, we found that after a single fraction of 8Gy, there was a delayed, but significant increase in intratumoral adenosine starting 48-hour post-treatment and persisting through 72-hour post-treatment (Figure 2A). To understand the implications of irradiation on myeloid and lymphoid cell populations, we conducted tumor immunophenotyping of EO771 tumors 1-, 3-, 5- and 11-days post-irradiation (8Gy X 3) by flow cytometry. In irradiated tumors, the proportion of CD11b+ cells was significantly higher than untreated tumors one day (day 1) after tumor irradiation, followed by a steady downward trend through day 11 (Figure 2B). Conversely, there was a significant decrease in the proportion of T cells, compared to untreated tumors on day 1, with recovery and a 2- and 6-fold increase 5- and 11-days post-irradiation, respectively (Figure 2C). These data illustrate that lymphocytes are most sensitive to irradiation in the acute setting (days 1-3), and with time, the abundance of T cells in TME recovers, as has been shown previously (17). Similar observations were made in the 4T1 model, with significant increase in adenosine (supplementary figure S2A) and lymphotoxicity observed at the acute phase post-irradiation (supplementary figure S2B, S2C). Deeper investigation of gene expression profiles by RNA-seq of CD45+CD11b+ cells sorted from tumors revealed an increase in expression of genes promoting expansion or infiltration of myeloid cells (*Ly6c2, CCR2, ITGAM, CSF3R, CSF1R, ADGRE1*) and genes suppressing anti-tumor responses *(TREM2, SIRPA, PPARG, ARG1)* on day 3 post-irradiation with a decline by day 5 (Figure 2D). In CD45+TCRβ+ T cells, at the early 3-day timepoint, there was decreased expression of activated, memory and effector T cells genes (*SELL*, *CD44*, *GZMA*), with increased expression of immune checkpoint genes (*Pdcd1, Ctla4*), exhaustion (*Tox*) and regulatory T cell genes (*FoxP3)* suggesting damage to T cell survival, activation and function due to irradiation (Figure 2E). Simultaneously, in sorted CD45- cells, on day 3, there was upregulation in expression of tissue remodeling genes (*KRT8*, *FBN1*, *COL3A1*) (Figure 2F). Overall, these results suggest that irradiation fosters myeloid cell induction and suppression of T cell function, thus driving immunosuppression in the TME acutely after treatment. Further, we found that radiation induced significant upregulation of A_2a_R on day 3 in CD45+CD11b+ myeloid cells (Figure 2G) and CD45+TCRβ+ T cells (Figure 2H) followed by a decline by day 5 and a simultaneous gradual increase in NT5E from days 1-5 in CD45- cells (Figure 2I). These data suggest a potential role for adenosine signaling in acute radioresistance.

**Figure 2:**
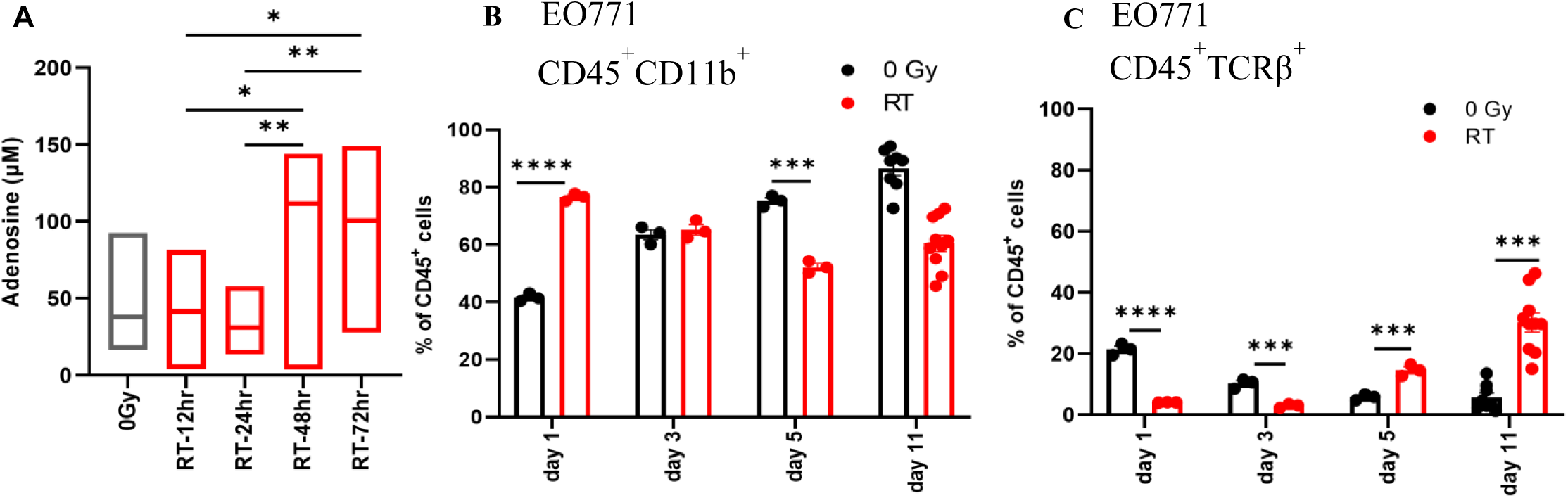
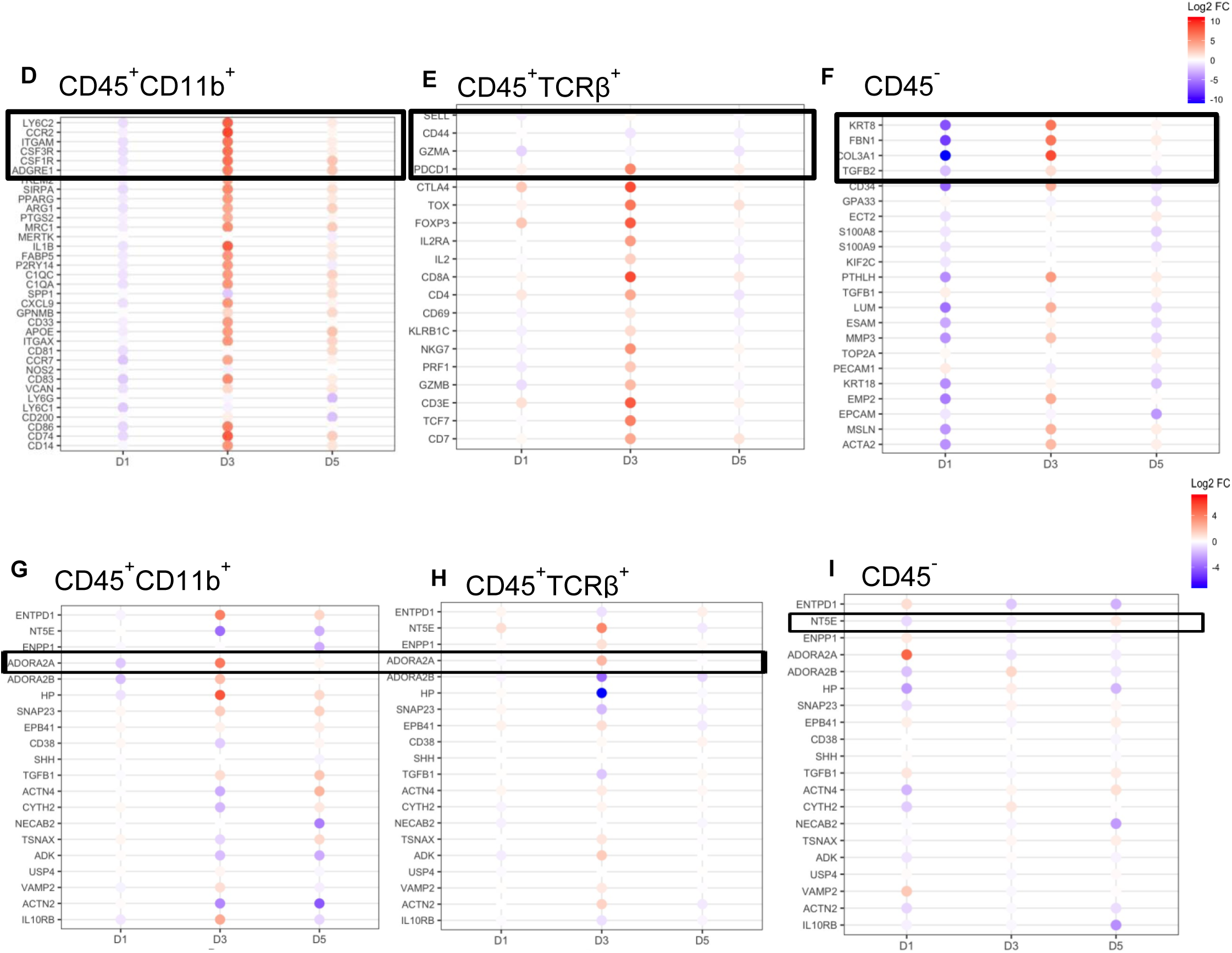
(A) Fold change in intra-tumoral adenosine concentrations in irradiated (8Gy x 1) EO771 tumors 12-, 24-, 48-, and 72-hours after treatment calculated with respect to adenosine concentration in unirradiated tumors. Abundance (Mean ± SEM) of (B) myeloid cells (CD45^+^CD11b^+^) and (C) T lymphocytes (CD45^+^TCRb^+^) in EO771 tumors on days 1,3,5,11 after irradiation (8Gy x 3) compared to unirradiated controls. Data are representative of two separate experiments, n=10-12 mice/group. (***) p<0.001, (****) p<0.0005. Log2 fold changes in gene expression in irradiated cells versus the corresponding unirradiated cells on day 1 (D1), 3 (D3), 5 (D5) in (D) CD45^+^CD11b^+^ cells (E) CD45^+^TCRb^+^ cells (F) CD45^-^ cells sorted from intra-tumoral cell suspensions (n=3 mice/group). Log2 fold changes in gene expression of adenosine signaling genes in irradiated cells versus the corresponding unirradiated cells on day 1, 3, 5 in (G) CD45^+^CD11b^+^ cells (H) CD45^+^TCRb^+^ cells (I) CD45^-^ cells sorted from intra-tumoral cell suspensions (n=3 mice/group).

### Loss of CD73 in cancer cells and A2aR in myeloid cells improves tumor radiosensitivity

Radiation increases suppressive adenosine in TME. We sought to determine the contribution of CD73 and ENPP1 to intratumoral adenosine during radiation. Loss of CD73 caused near-complete loss of adenosine production from CD73^-/-^ EO771 cells confirming that indeed CD73 is required for adenosine production (Figure 3A). Loss of ENPP1 caused a small but significant 1.2-fold decrease in adenosine concentration (Figure 3A) demonstrating that ENPP1 is not a major source of AMP, the substrate for adenosine production by CD73. Irradiation of the EO771 cells caused a 1.2-fold increase in extracellular adenosine (Figure 3A). Notably, loss of ENPP1 mitigated the radiation-driven increase in adenosine, suggesting that radiation-induced cGAS-mediated cGAMP production at least partially contributes to radiation-driven increase in adenosine (Figure 3A).

**Figure 3:**
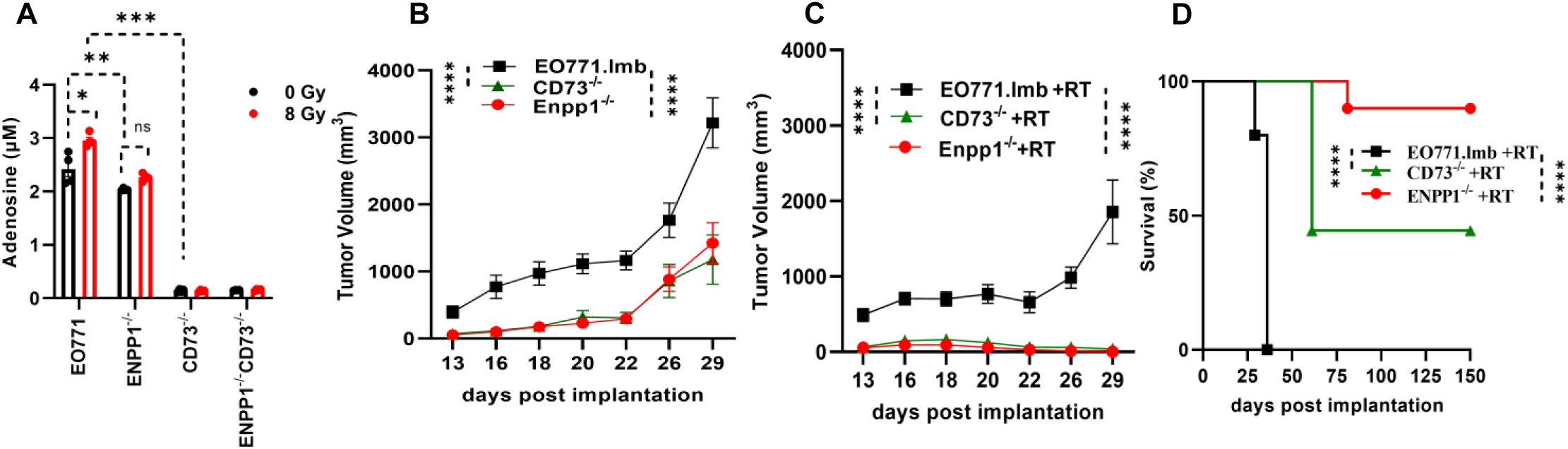
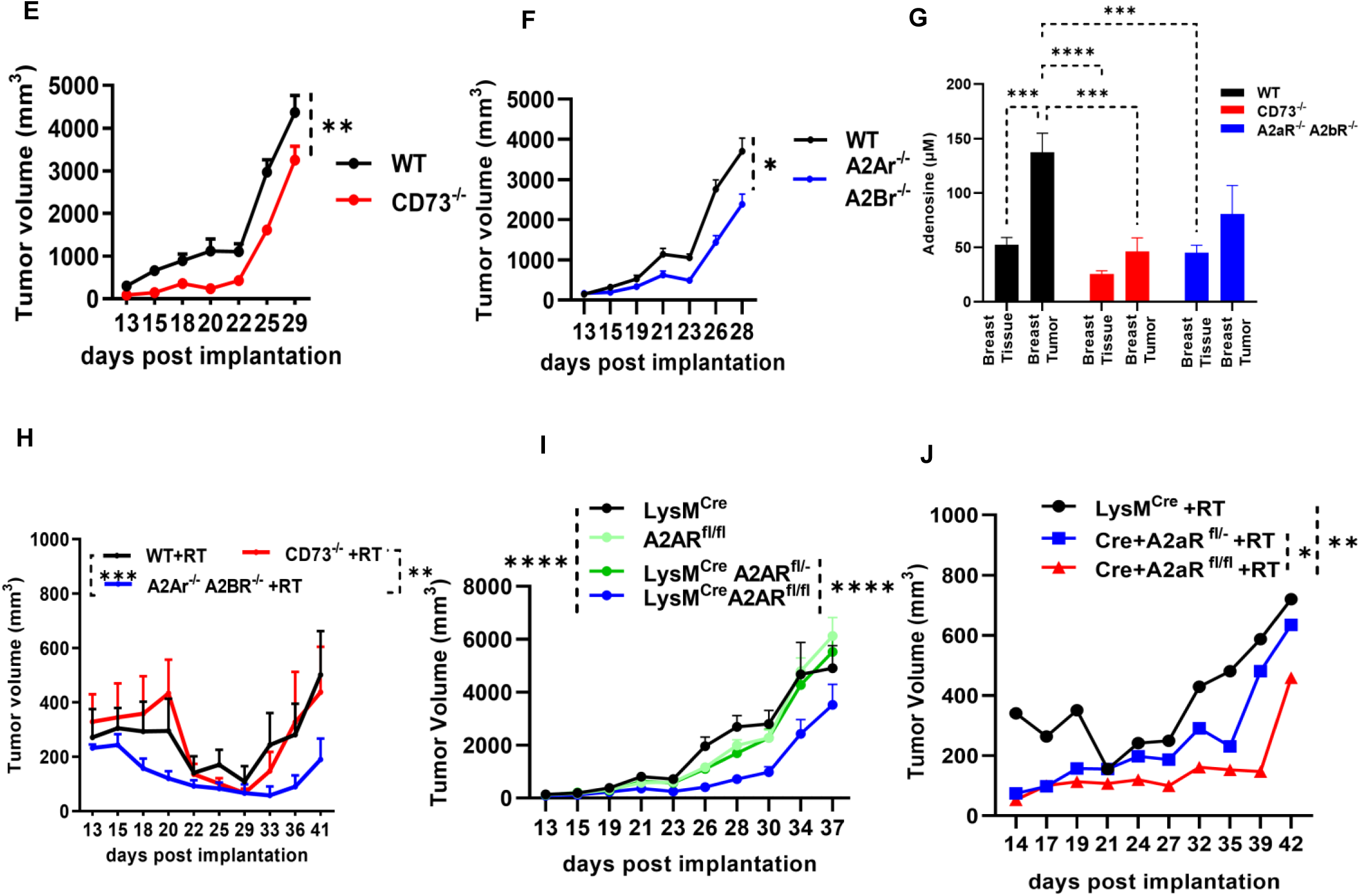
(A) Adenosine concentration (Mean ± SEM) measured in cell-free supernatant from WT (EO771.lmb), ENPP1^-/-^, CD73^-/-^, ENPP1^-/-^CD73^-/-^ cells grown *in vitro,* 18 hours after irradiation (8 Gy x 1), compared to untreated controls. (B) Tumor volume (Mean ± SEM) from WT mice bearing WT EO771, CD73^-/-^ EO771, or Enpp1^-/-^ EO771 tumors. (C, D) Tumor volume (Mean ± SEM) and survival of WT mice bearing WT EO771, CD73^-/-^ EO771, or Enpp1^-/-^ EO771 tumors irradiated (8Gy x 3) on days 13, 14, 15 post-implantation. (E, F) Tumor volume (Mean ± SEM) of orthotopic EO771 tumors implanted in wildtype (WT, C57BL6/J), CD73^-/-^ and A_2a_R^-/-^A_2b_R^-/-^ mice. (G) Intra-tumoral adenosine concentration (Mean ± SEM) in paired normal murine breast tissue and EO771 tumors implanted in WT, CD73^-/-^ and A_2a_R^-/-^A_2b_R^-/-^ mice. (H) Tumor volumes (Mean ± SEM) from WT (C57BL6/J), CD73^-/-^, A_2a_R^-/-^A_2b_R^-/-^ mice implanted with EO771 cells irradiated (8 Gy x 3) on days 13, 14, 15 post-implantation. Tumor volumes (Mean ± SEM) from (I) untreated and (J) radiation-treated (8Gy x 3) tumors from LysM^Cre^, A_2a_R^fl/fl^, LysM^Cre^A_2a_R^fl/-^, LysM^Cre^A_2a_R^fl/fl^ mice implanted with EO771 cells. Data are representative of two separate experiments, n=10-12 mice/group (*) p<0.05 (**), p<0.01(***), p<0.001, (****)p<0.0005.

Our data suggests that adenosine signaling is a cooperative system driven by both cancer and immune cells. Striking expression of CD73 on cancer cells and A_2a_R on immune cells in human breast cancer specimens (Figure 1D, Figure 1E) and murine breast tumors (Figure 1G, 1I) fueled our interest in investigating their roles in contributing to immune-mediated radioresistance in murine breast tumors. Knockdown of CD73 or ENPP1 in tumor cells improved tumor control to a similar degree (Figure 3B), suggesting the importance of tumor cell-derived adenosine from CD73 and the cGAMP-STING pathway activation via loss of ENPP1 to tumor progression in the absence of radiotherapy. However, after tumor irradiation (8Gy x 3), WT EO771 tumors regressed up to 10 days post-RT after which 100% regrew requiring euthanasia 15 days post-RT. Conversely, ENPP1^-/-^ and CD73^-/-^ EO771 tumors, demonstrated a significantly more durable response to irradiation whereby 90% of mice bearing ENPP1^-/-^ tumors and 44% of mice bearing CD73^-/-^ tumors were tumor-free on day 20 post-tumor irradiation (Figure 3C, 3D). These data suggest that loss of adenosine-producing CD73 and cGAMP-metabolizing ENPP1 from cancer cells alone is sufficient to improve radioresponsivness, illustrating the powerful contribution of tumor cell-derived adenosine and the cGAS-STING pathway to oncologic outcomes.

To investigate the contribution of host CD73 and adenosine receptor (A_2a_R and A_2b_R) expression to radioresponsivness, wildtype EO771 tumors were grown in CD73^-/-^ and A_2a_R^-/-^A_2b_R^-/-^ mice. In untreated mice, tumor control was improved in CD73^-/-^ mice (Figure 3E) and A_2a_R^-/-^A_2b_R^-/-^ mice (Figure 3F). Delay in tumor growth in CD73^-/-^ mice correlated with significantly lower intra-tumoral adenosine, compared to WT mice. Whereas there was no significant increase in intra-tumoral adenosine in the A_2a_R^-/-^A_2b_R^-/-^ mice (Figure 3G). In irradiated mice, loss of CD73 from tumor-bearing mice was insufficient to improve tumor control but loss of A_2A_R and A_2B_R from the mice did improve radioresponsivness (Figure 3H). This suggests that in the context of tumor irradiation, the contribution of host derived CD73 is insufficient to influence adenosine levels that offer a clinical influence on radioresponse. Whereas, lack of host derived A_2a_R and A_2b_R mediated adenosine signaling offers substantial improvement in tumor radiosensitivity.

In the EO771 model, the intratumoral myeloid compartment expresses higher levels of the adenosine receptor, A_2a_R, compared to lymphocytes, including T cells (Figure 1I). We posited that the myeloid compartment may indeed be a major mediator of the suppressive action of adenosine. To investigate this, we generated mice that lack expression of one or two alleles of the A_2a_R in myeloid cells (LysM^Cre^). Compared with A_2a_R^fl/fl^ and LysM^cre^ controls, mice deficient in both alleles of A_2a_R in myeloid cells (LysM^Cre^A_2a_R^fl/fl^) exhibited significant delay in tumor progression and a 3-fold decrease in tumor volume 30 days post-implantation that persisted beyond day 30 (Figure 3I). However, when tumors were irradiated, the loss of A_2a_R expression in the myeloid cells resulted in a 4-fold decrease in tumor volume compared to control (Figure 3J). Spider plots and the slope of tumor re-growth curves reveal exponential tumor growth in 83% LysM^Cre^ control mice (5 of 6 mice) (supplementary figure S3A, S3C, S3D) while only 50% of tumors grew exponentially in LysM^Cre^A_2a_R^fl/fl^ mice (4 of 8 mice) (supplementary figure S3B, S3C, S3E). Taken together, these data highlight the important role of adenosine signaling in myeloid cells as a mediator of radioresponsivness.

### Adenosine blockade improves tumor control by radiotherapy and checkpoint inhibition in EO771 tumor bearing mice

Radiotherapy offers tumor control but drives an immunosuppressive TME (3). At the same time, it is established that EO771 murine tumors have a modest response to anti-PD-1 (20). We postulated that blockade of CD73 and A_2a_R may partially alleviate the immunosuppressive environment in the tumor and improve response to radiotherapy and anti-PD-1. Leveraging our EO771 and 4T1 breast cancer models, we conducted therapeutic trial. Mice were treated with adenosine signaling modulators, AB680 (CD73 inhibitor) and AB928 (dual antagonist of A_2a_R/A_2b_R), from day 8 – 24 post implantation (Figure 4A). When tumors reached ∼100-150 mm^3^ (day 13), they were irradiated (8Gy x 3 daily). The day after tumor irradiation, anti-PD-1 (AB122) was started and delivered every 2 days for three doses (Figure 4A). In EO771 model, in the absence of tumor irradiation, the combined immunotherapies, AB680, AB928 and anti-PD-1, had no significant effect on tumor growth compared to vehicle treated mice (Figure 4B). Radiotherapy alone caused a significant decrease in tumor growth that was further improved with the addition of adjuvant anti-PD-1 (Figure 4B, 4C). The combination of adenosine inhibitors and radiotherapy improved tumor control compared to radiotherapy alone or radiotherapy with anti-PD-1 (Figure 4B, 4C). The addition of adenosine inhibitors to radiotherapy and anti-PD-1 improved the overall response rate, with 7/10 mice having no palpable evidence of tumor compared to 2/9 mice treated with adenosine inhibitors and radiotherapy by day 25 (Figure 4D). With longer follow-up (day 32-42) tumor regrowth was slowest in the combined therapy group: adenosine inhibitors, anti-PD-1 and radiotherapy (Figure 4C, Figure 4E). Similarly, a therapeutic trial in the murine 4T1 TNBC model demonstrated significantly improved tumor control after triple therapy comprising adenosine pathway inhibitors, radiotherapy and anti-PD-1 compared with either adenosine inhibition plus radiotherapy or radiotherapy plus anti-PD-1 (Figure 4F). Collectively, findings across both breast cancer models indicate that dual targeting of CD73 and A2A receptor enhances responses to radioimmunotherapy and results in superior tumor control.

**Figure 4:**
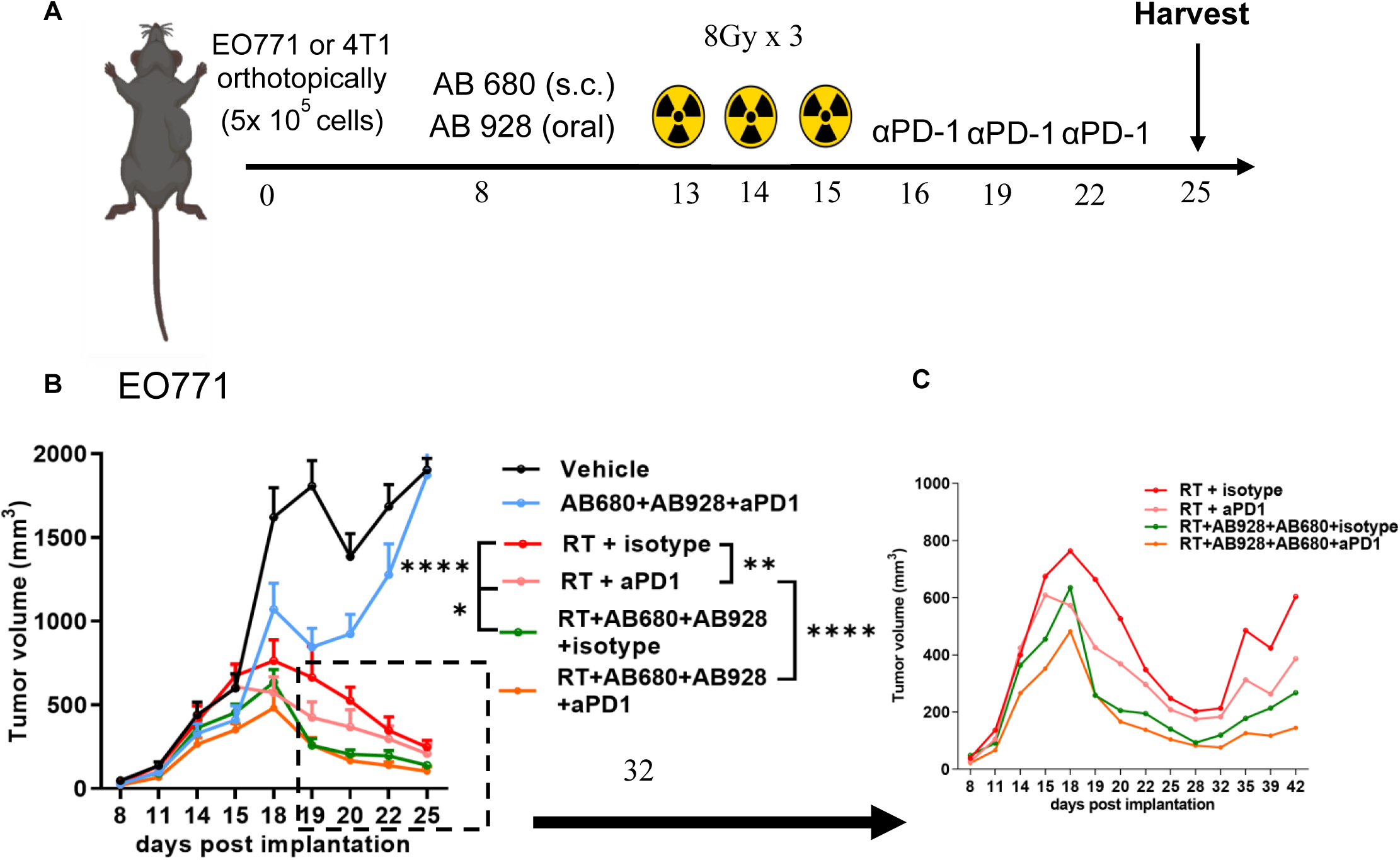
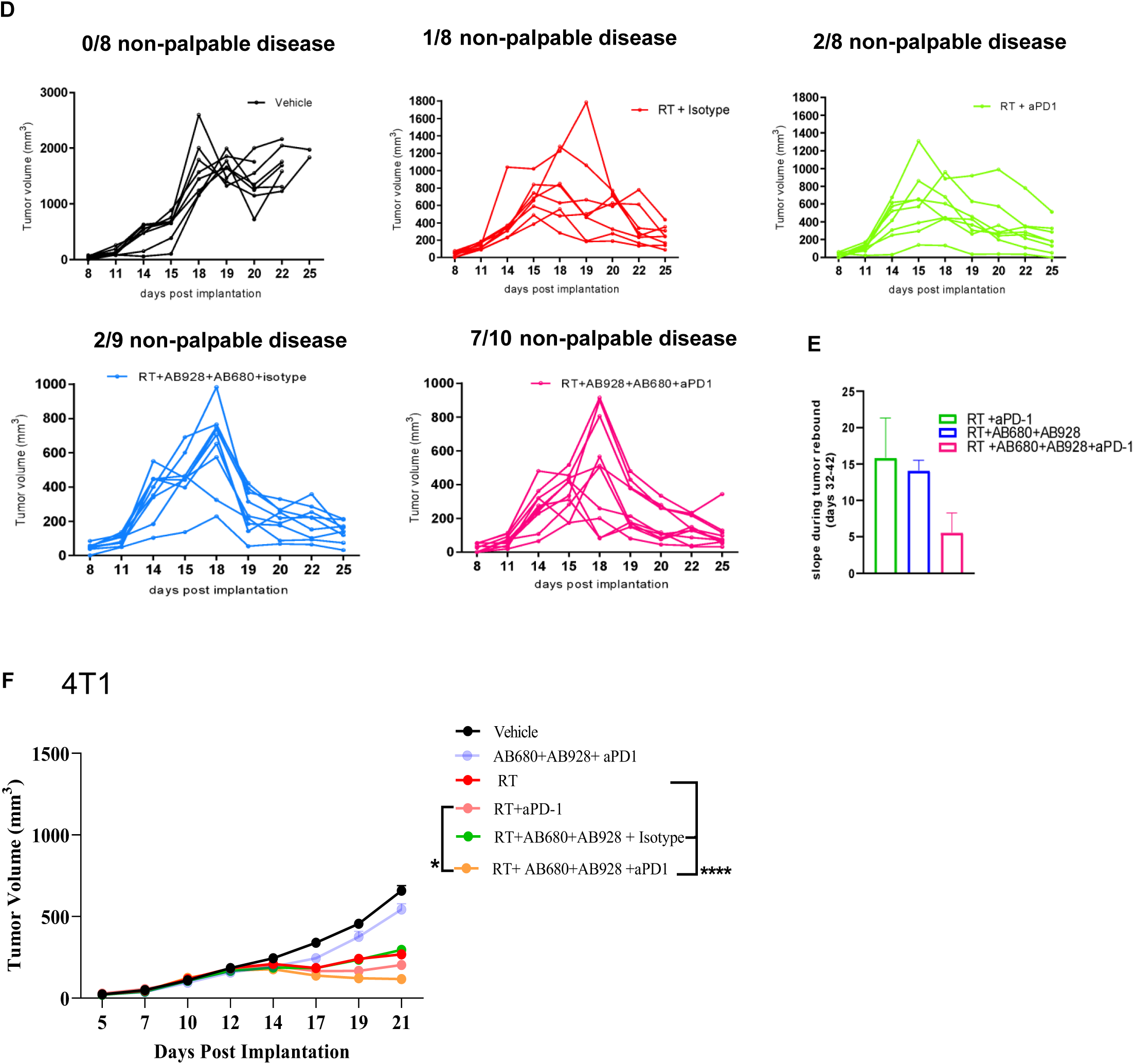
(A) Schematic summary of treatment schedule for EO771 or 4T1 orthotopic breast cancer model. (B) Tumor volume measurements (Mean ± SEM) beginning 8 days after EO771 implantation (n=8-10 mice/group). AB680 (quemliclustat, CD73 inhibitor), AB928 (etrumadenant, A_2a_R/A_2b_R antagonist), αPD1 (zimberelimab, anti-PD-1 antibody), isotype control (IgG2 antibody), tumor irradiation (RT, 8 Gy x 3; 12-, 13-, 14- days after implantation). (C) Tumor volumes from days 8-42 of irradiated mice treated in combination with isotype control, anti-PD-1, AB928 and AB680, AB928 and AB680 and anti-PD-1. (D) Spider plots of tumors from mice belonging to all treatment groups implanted with EO771 cells. (E) Slopes (Mean ± SEM) during tumor rebound period (days 32-42) from growth curves of mice belonging to irradiation and anti-PD-1, irradiation and AB680 and AB928, irradiation and AB680 and AB928 and anti-PD-1 treatment groups. (F) Tumor volume measurements (Mean ± SEM) beginning 8 days after 4T1 implantation. Data are representative of two separate experiments, n=8-10 mice/group (*) p<0.05 (**), p<0.01, (****) p<0.0005.

### Combination treatment with adenosine blockade, radiotherapy and checkpoint inhibition promotes M1 TAMs and improves effector function of T cells in EO771 tumors

In addition to improving tumor control, we hypothesized that blockade of the adenosine signaling axis will unmask the immunogenic potential of radiation by restoring cytotoxic T cell function thus promoting anti-tumor immunity. To visualize global changes after different treatments, we performed t-stochastic neighbor embedding (tSNE) analysis on 50,000 concatenated CD45+ immune cells from each sample collected 11 days after tumor irradiation. Through marker visualization and gating validation, we identified four phenotypically distinct clusters in the tSNE plot. Clusters of lymphoid cells were localized predominantly in center and on left whereas myeloid cells were clustered on the top and right (Figure 5A). Comparative analysis demonstrated an increase in the relative abundance of lymphoid cell clusters post radiotherapy, adenosine blockade and anti-PD-1 compared to vehicle or adenosine blockade treated mice (Figure 5B). Additionally, compared to all other groups, there was a shift in the spatial distribution of cells in myeloid clusters suggesting alterations in transcriptional state following combination therapy (Figure 5B). Deeper immunophenotyping by manual gating revealed that in mice receiving treated with the combination of radiotherapy, adenosine blockade and anti-PD-1, there was a shift in the macrophage compartment towards anti-tumorigenic, F4/80^+^MHCII^+^ (M1-like) TAMs from pro-tumorigenic F4/80^+^MHCII^-^ (M2-like) TAMs and decrease in the proportions of intra-tumoral PMNs and monocytes compared to vehicle treated mice (Figure 5C). These data suggest that combination therapy can reprogram the myeloid compartment from a suppressive myeloid cell milieu towards a more immune permissive TME. In lymphoid compartment, we observed an increase in the proportions of intratumoral CD4 T cells, CD8 T cells, proliferating CD8 T cells and the CD8:CD4 ratio in mice treated with radiotherapy alone or in combination with anti-PD-1. The addition of adenosine inhibitors to RT and anti-PD-1 did not further influence the abundance of these key players of adaptive immunity (Figure 5D). EO771 tumors consist of minimal proportions of DCs (0.5%), NK cells (3%), B cells (0.9%) compared to myeloid and lymphoid cells that did not change significantly with treatment (supplementary Figure S4). While, the combination of radiotherapy, adenosine blockade and anti-PD-1 did not modulate proportions of CD4 T cells, CD8 T cells, proliferating CD8 T cells in the TME, we hypothesize that T cell effector cytokines may indeed be increased following the combination treatment.

**Figure 5:**
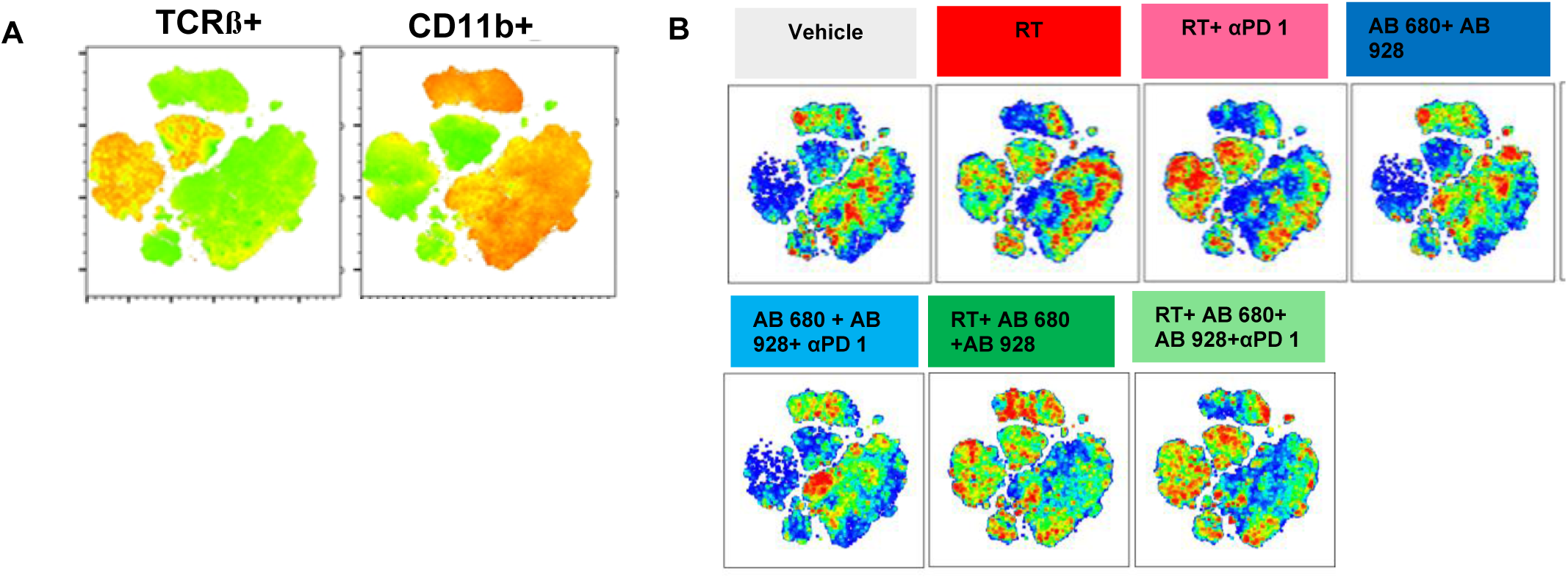
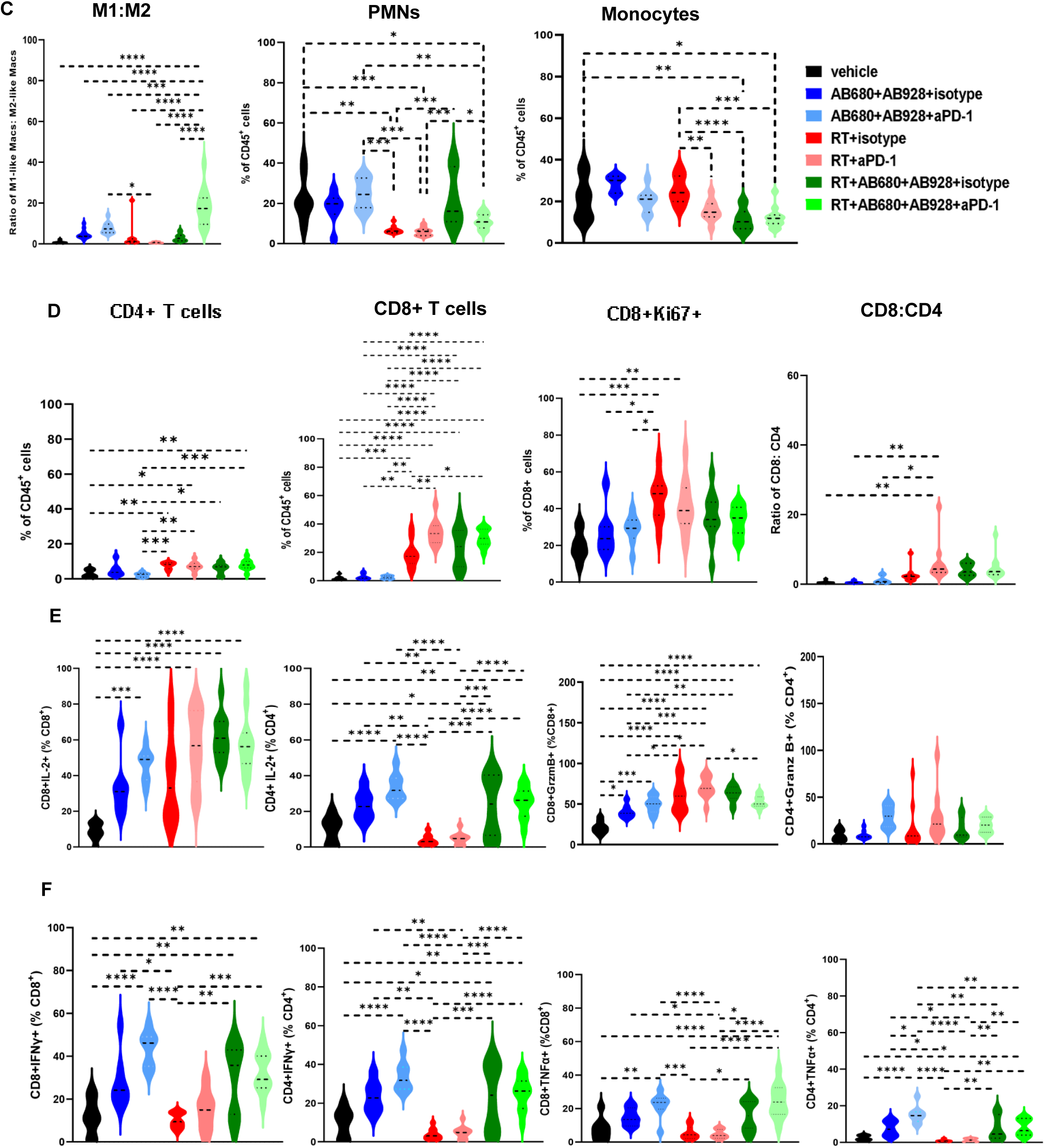
(A) TCRß+ and CD11b+ clusters from CD45+ immune cells in tSNE plots (B) tSNE plots of CD45+ immune cells from mice belonging to different treatment groups (C) Ratio of M1-like: M2-like macs (Mean ± SEM), proportion of immune compartment (CD45+) that are PMNs (Mean ± SEM), Mo (Mean ± SEM), 11 days after radiation in every treatment group. (D) Proportion of immune compartment (CD45+) that are CD4+ T cells (Mean ± SEM), CD8+ T cells (Mean ± SEM), proliferating (Ki67+) CD8+ T cells (Mean ± SEM) and ratio of CD8+: CD4+ T cells (Mean ± SEM). Proportion of CD8 T cells and CD4 T cells that express (E) IL-2, granzyme B (Mean ± SEM), (F) IFNγ, TNFα (Mean ± SEM) per treatment group. Data are representative of two separate experiments, n=10-12 mice per group. (*) p<0.05, (**) p<0.01, (***) p<0.001, (****) p<0.0005. M1-like macs (CD45^+^CD11b^+^Ly6C^-^F4/80^+^MHCII^+^), M2-like macs (CD45^+^CD11b^+^Ly6C^-^F4/80^+^MHCII^-^), PMN–neutrophil (CD45^+^CD11b^+^Ly6G^+^), Mo–monocytes (CD45^+^CD11b^+^Ly6C^+^). Data are representative of two separate experiments, n=10-12 mice/group. (*) p<0.05 (**), p<0.01(***), p<0.001, (****) p<0.0005.

To test this hypothesis, we conducted an *ex vivo* stimulation assay to quantify the expression of effector cytokines by intratumoral T cells collected after treatment. Radiotherapy alone increased the production of IL-2 and granzyme B in CD8 T cells (Figure 5E) but did not alter expression of IFN γ and TNFα by CD8 or CD4 T cells (Figure 5F). Adenosine blockade alone did not alter T cell abundance (Figure 5D) but did significantly increase the expression of effector cytokines including IL-2, IFN γ and TNFα by CD8 and CD4 T cells (Figure 5E, 5F) and granzyme B by CD8 T cells (Figure 5E). When adenosine blockade was combined with radiotherapy, effector cytokine production was rescued for both CD8 and CD4 T cells, increasing the expression of IFN γ and TNFα by CD8 and CD4 T cells and IL-2 by CD4 T cells (Figure 5E, 5F). These results demonstrate that the addition of adenosine inhibitors and anti-PD-1 to radiotherapy improves effector T cell function and tumor control, whereas adenosine inhibitor and anti-PD-1 improves effector cytokine production by T cells without tumor control benefit.

### Tumor, immune and adenosine gene signatures predict response to radio-immunotherapy in human breast carcinoma

To assess the translational potential of adenosine signaling inhibition and radiotherapy in breast cancer, we analyzed a publicly available single-cell RNA sequencing dataset generated from biopsies collected from women with localized ER-PR-HER2- breast cancer (TNBC) at (1) baseline, (2) 3-weeks after neoadjuvant anti-PD-1 monotherapy, and (3) 3-weeks after neoadjuvant breast irradiation (8Gy x 3) and anti-PD-1 therapy before proceeding with standard neoadjuvant chemotherapy (26). We analyzed data from three patients who had complete pathologic responses (pCR, ‘responders’) and three ‘non-responders’ (no pCR) (supplementary table 5). Just 3-weeks after a non-curative dose of radiation (8 Gy x 3) plus anti-PD-1, and before exposure to neoadjuvant chemotherapy, the responders had a significant reduction in malignant cells (denoted by copy number variation (CNV) high myoepithelial and myoepithelial/ductal cells (supplementary Figure S5a), compared to pre-treated or anti-PD-1 treated tissue whereas malignant cells persisted in similar abundance in the non-responders. These data raise the hypothesis that the pathologic response to neoadjuvant anti-PD-1 and radiotherapy may serve as a predictive biomarker and guide patient selection for intensive chemotherapy regimens (Figure 6A). Within the CD45+ compartment of tumor biopsies, we found no significant change in the abundance of myeloid cell types over the course of treatment (supplementary Figure S5b). However, there was a decrease in regulatory T cells (Tregs) and exhausted, antigen-reactive CD8 T cells (CD8+ T cells ExRe) in the responders after anti-PD-1 alone that further decreased after anti-PD-1/radiotherapy. In the non-responders, however, we observed an increase in Tregs and CD8+ T cells ExRe after anti-PD-1 monotherapy that were sustained or increased following the combination of anti-PD-1/radiotherapy (Figure 6B). These data suggest that suppressive immune cells in the TME may predict response to therapy.

**Figure 6:**
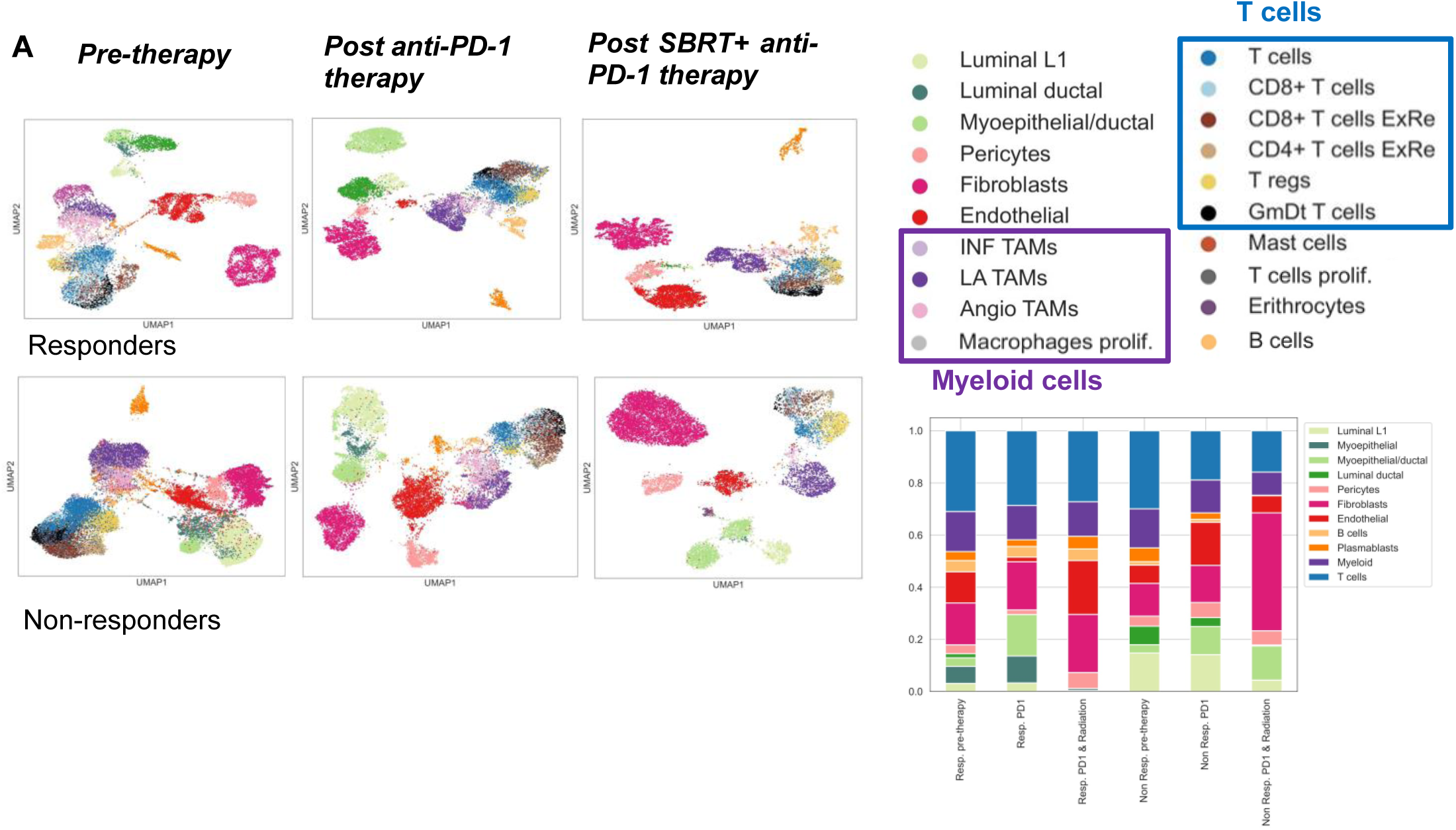
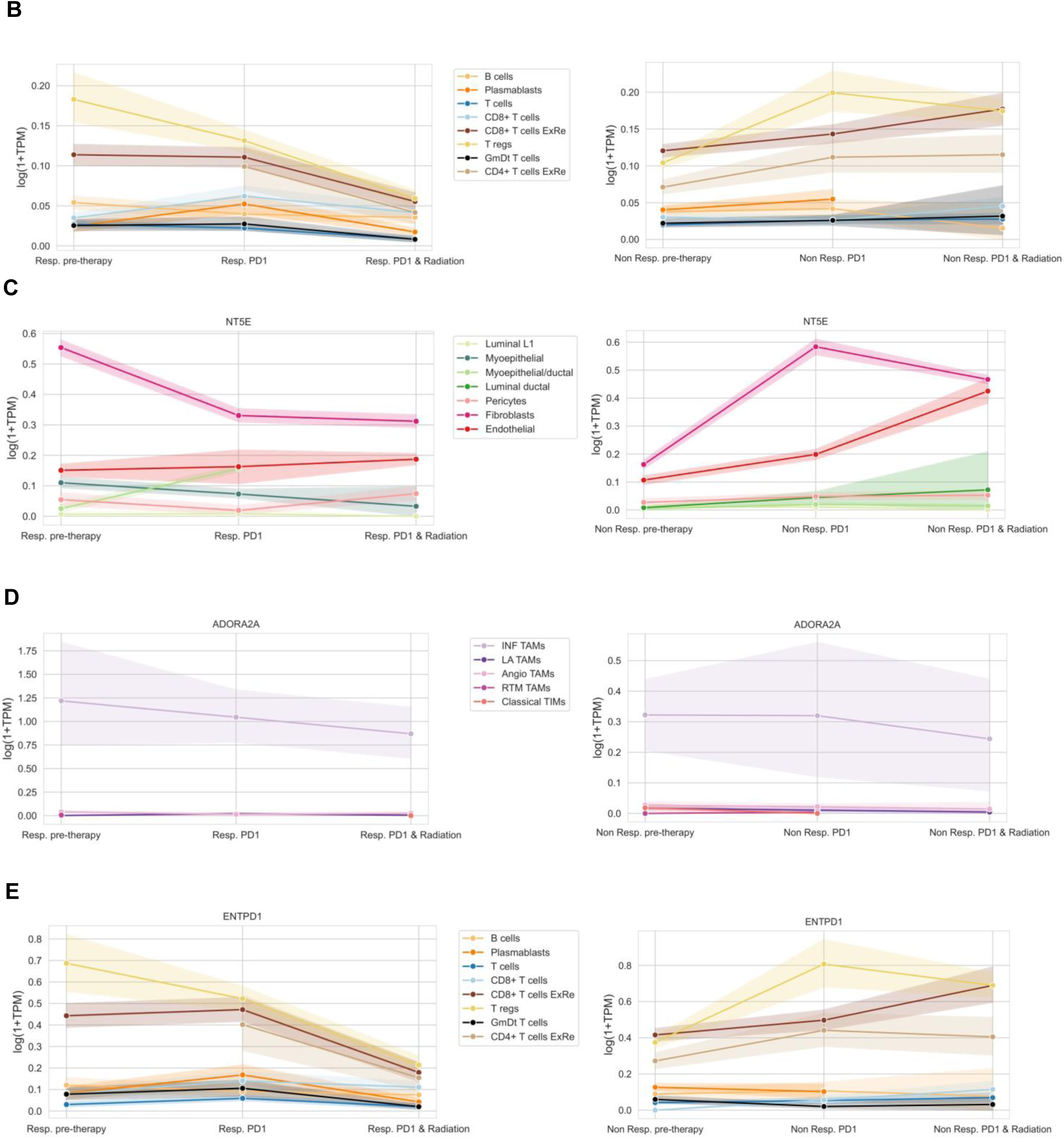
(A) Leiden clustering of non-immune and immune cell populations from scRNA seq data from 3 responder and 3 non-responder patients collected at pre-therapy, post anti-PD-1 therapy and post-SBRT plus anti-PD-1 therapy. (B) Abundance of lymphoid cells in responders and non-responders at different stages of therapy. Proportions of (C) non-immune cells expressing NT5E (CD73) in responders and non-responders at different stages of therapy. (D) myeloid cells expressing A_2a_R in responders and non-responders at different stages of therapy (E) lymphoid cells expressing ENTPD1 (CD39) in responders and non-responders at different stages of therapy.

Inspection of adenosine signaling genes in individual populations yielded insight about their potential role in anti-PD-1 and radiotherapy response in patients. Baseline expression of *NT5E* (CD73) in tumor fibroblasts and *A2aR* on inflammatory tumor-associated macrophages (INF TAMs, LAMP3+) was high, corroborating our findings in our pre-clinical murine breast cancer model. CD73 expression decreased in fibroblasts after combined radiation and anti-PD-1 therapy in responders whereas, in non-responders, its expression increased in both fibroblasts and endothelial cells in response to therapy (Figure 6C). A_2a_R expression remained unchanged after neoadjuvant anti-PD-1 or combination therapy (Figure 6D). *ENTPD1* (CD39) expression was highest in the immune compartment. Its expression significantly decreased in Tregs, ExReCD8 and CD4 T cells after anti-PD-1/radiotherapy in responders, whereas *ENTPD1* expression increased in the non-responders in these cells (Figure 6E). Based on this small cohort of patients with TNBC, these data suggest that changes in CD73 and CD39 in the setting of radioimmunotherapy may be prognostic in breast cancer patients and warrant further investigation of the adenosine signaling axis in this context.

## DISCUSSION

Radiotherapy remains a cornerstone of cancer treatment, yet durable local control is frequently undermined by adaptive resistance mechanisms within the TME. Herein, we show that at baseline, breast tumors are rich in suppressive adenosine compared to non-malignant tissue (Figure 1A,1B) and indeed tumor irradiation increases adenosine in the TME (Figure 2A). While adenosine has long been recognized as an immunosuppressive metabolite, our data extends this paradigm by demonstrating that both tumor and immune compartments actively contribute to adenosine accumulation and sensing following irradiation, creating a self-reinforcing suppressive niche. Across multiple cancer types, it has been shown that expression of CD73 by tumor cells and A_2A_R, in the immune compartment of tumors is prognostic of worse outcomes (21, 22). Indeed, we confirmed that among the heterogeneous breast carcinomas of mice (EO771 and 4T1) and humans, CD73 is most highly expressed by tumor cells themselves (Figure 1D, 1F, 1H) while A_2A_R is more highly expressed in the immune compartment (Figure 1E), especially by intratumoral myeloid cells (Figure 1I). Breast carcinomas are frequently enriched in immunosuppressive myeloid populations (27). In this context, we propose that distinct components of the TME coordinate to regulate the adenosine signaling axis, with tumor cells being a major metabolic source of adenosine, myeloid cells serving as an adenosine-driven suppressive amplifier, and effector lymphocytes being subject to suppression. In the absence of Enpp1 or CD73 expressed by tumor cells, adenosine production is impaired (Figure 3A) and knock out of Enpp1, increases the abundance of cGAMP for STING pathway activation resulting in improved tumor control and overall survival (Figure 3C, 3D). However, decreased production of adenosine by the non-tumor elements of the TME (CD73^-/-^ mice) during radiation did not influence tumor control (Figure 3H). This illustrates that the tumor cell-intrinsic contribution to adenosine influences radioresponsiveness of the tumor, but adenosine produced by non-tumor cells in the TME does not. Consistent with our observations, Wennerberg et al (13) reported that CD73 expression on cancer cells is the critical determinant limiting the response to radiotherapy. Though expressed at very low levels, loss of A_2a_R/A_2b_R in the TME (figure 3F, 3H) and loss of A_2a_R specifically from the myeloid compartment (LysM^Cre^-mediated deletion) significantly improved tumor control of untreated tumors (Figure 3I), as shown previously (23), and also during tumor irradiation (Figure 3J). These data highlight the critical role of adenosine signaling axis in the myeloid compartment maintaining a suppressive, tumor-permissive TME. This is corroborated by high expression of A_2a_R by TAMs in EO771 and PMNs in 4T1 breast carcinoma, suggesting that A_2a_R signaling in these myeloid cell subsets contributes to radioresistance and tumor progression in both models. To our knowledge, this is the first study to demonstrate that adenosine production by CD73 expressed by tumor cells and engagement by the A_2a_R-expressing myeloid cells promotes immune-mediated radioresistance in breast cancer. Tumor irradiation converts tumors into in situ vaccines through immunogenic cell death and antigen release and promotes adaptive anti-tumor immune responses (24, 25). Simultaneously, radiotherapy reinforces suppressive immune changes including myeloid and regulatory T cells (28) that counter the therapeutic benefit of radiation. We show that, that in the acute setting (up to 3 days) radiotherapy induces profound metabolic and inflammatory perturbations in TME with lymphotoxicity, impaired cytotoxic T cell responses, upregulation of anti-tumor suppressing transcriptional programs and adenosine signaling in parallel (Figure 2). From a translational perspective, these findings have direct therapeutic implications. Our data demonstrate that, combining CD73 and A_2a_R/A_2b_R antagonist during and after tumor irradiation to suppress the adenosine signaling axis followed by anti-PD-1 therapy (Figure 4A), improves therapeutic benefit of radiation (Figure 4B, 4C, 4D). Over time, as the effect of tumor irradiation subsides, lymphocytes recover and become significantly more abundant with improved CD8:CD4 T cell though with limited effector function. However, therapeutic targeting of adenosine signaling axis augments T cell effector responses (Figure 5E, 5F). Clinical trials combining radiotherapy with immune checkpoint blockade alone have yielded mixed results, highlighting the need to identify dominant immunosuppressive pathways that limit synergy. Our data supports the rationale for simultaneous disruption of adenosine signaling in both tumor and immune compartments to achieve meaningful radiosensitization, overcoming the limited efficacy observed with monotherapies targeting individual components of this pathway.

Furthermore, components in the adenosine pathway may serve as predictive biomarkers of radiotherapy response. From our investigation of scRNAseq data generated by Shiao et al (26), we confirmed expression of CD73, A_2a_R, CD39 in the TME of human TNBC tumors that differed in expression by responders and non-responders (Figure 6C, 6D, 6E). Although these data are derived from a small cohort of patients, CD39 and CD73 may serve as prognostic and predictive biomarkers of response to radiation and thus more likely to benefit from therapy intensification or incorporation of adenosine blockade. Given the availability of clinically advanced agents targeting this pathway, immediate translation is possible.

There are multiple ongoing studies testing the clinical efficacy of the combination of adenosine signaling modulators and radiation for patients with cancer. Among these, AIRPanc (NCT06048484) study is testing the neoadjuvant CD73 (AB680), A_2a_R/A_2b_R (AB928) and anti-PD-1 (AB122) in combination with SBRT for locally advanced pancreatic cancer and the PANTHER study (NCT05024097) is testing anti-PD-1 (AB122) and A_2a_R/A_2b_R antagonist (AB928) with chemotherapy after short-course radiation for rectal cancer (29). For patients with metastatic lung cancer, A_2a_R inhibitor (JNJ-86974680) is combined with anti-PD-1 (cetrelimab) and radiation (NCT06116786). By linking radiation-induced metabolic stress to suppression of antitumor immunity, we define a mechanistic framework that explains therapeutic failure and reveals a tractable vulnerability. Integrating radiotherapy with adenosine-targeted interventions represents a promising strategy to unlock the full immunomodulatory potential of radiation and improve clinical outcomes.

## ACKNOWLEDGEMENT

Dr. Matt Walters from Arcus Biosciences kindly provided us AB928, AB680 and AB122. We thank Dr. Renu Nandakumar for adenosine estimation by LC-MS, Hongyan Tang for helping with IHC staining, Michael Kissner and Christopher Wu for support with FACS sorting and the Herbert Irving Comprehensive Cancer Center Molecular Pathology and Human Immune Monitoring Core Shared Resource, and Cancer Biostatistics Shared Resource.

