## Supplementary material for "Targeting CD73-A_2a_R-Mediated Adenosine Signaling at the Tumor-Immune Interface Overcomes Radioresistance": Supplemetary material

**Graphical Abstract**

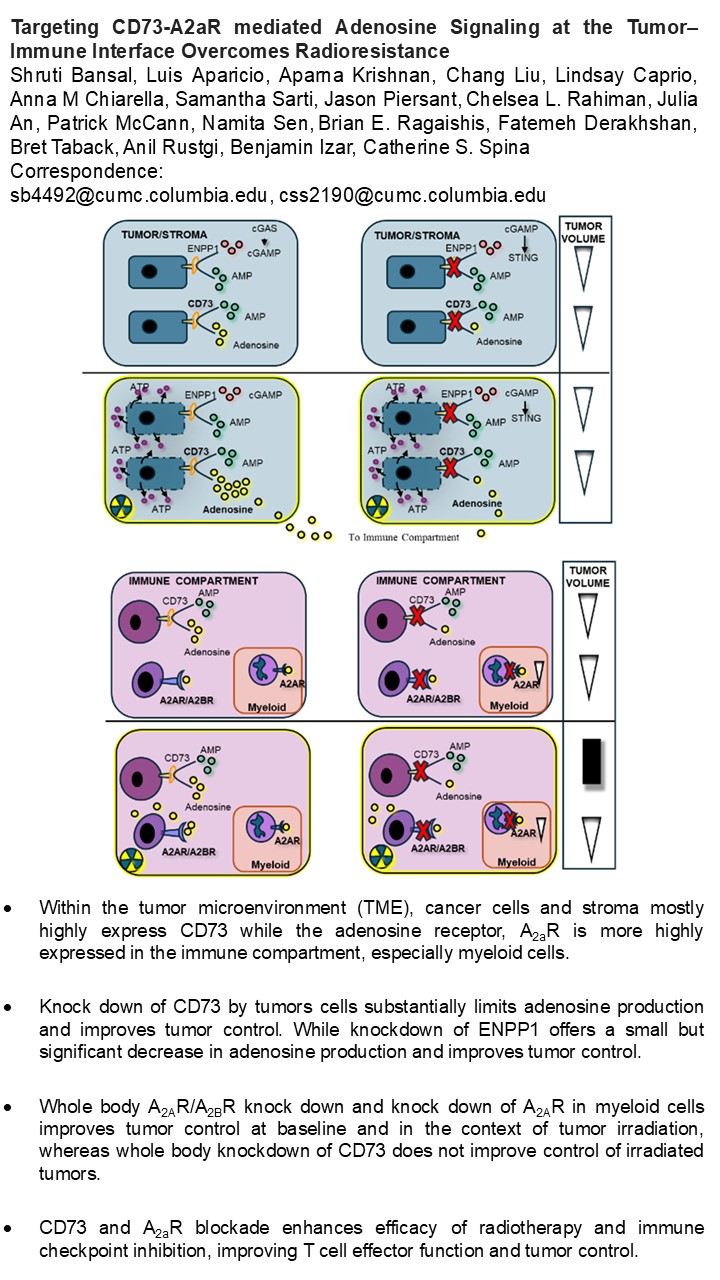

**Supplementary Results**

Figure S1: Frequency of neutrophils (PMNs, CD45^+^CD11b^+^Ly6g^+^), Monocytes (Mo, CD45^+^CD11b^+^Ly6c^+^) and Macrophages (Macs, CD45^+^CD11b^+^F4-80^+^Ly6G^-^Ly6C^-^) (Mean ± SEM) in (A) EO771 and (B) 4T1 tumors by spectral flow cytometry. Expression of adenosine signaling proteins on PMNs, Mo, Macs (Mean ± SEM) in (C) EO771 and (D) 4T1 tumors quantified using spectral flow cytometry

(A) EO771 CD45^+^ tumor infiltrate	(B) 4T1 CD45^+^ tumor infiltrate

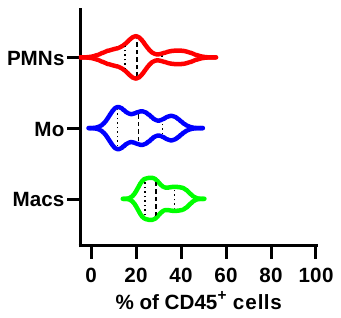

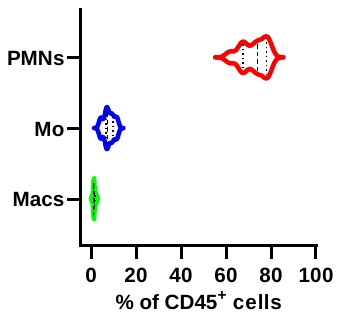

(C) EO771 CD45^+^ tumor infiltrate	(D) 4T1 CD45^+^ tumor infiltrate

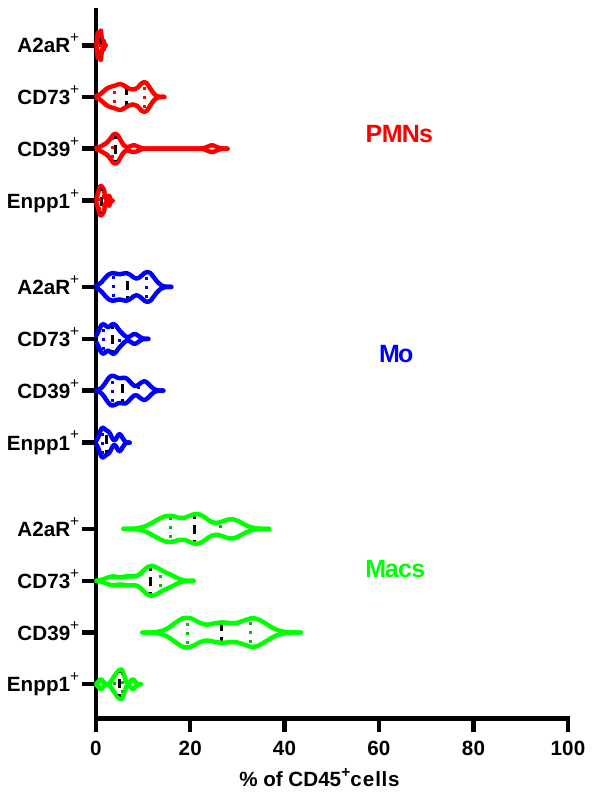

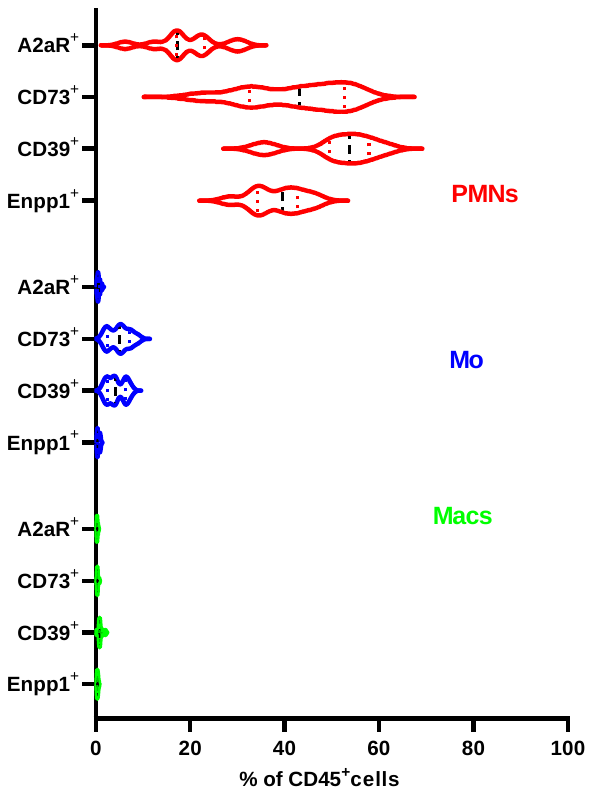

Figure S2: Fold change in intra-tumoral adenosine concentrations in irradiated (8Gy x 1) 4T1 tumors 16-hours after treatment calculated with respect to adenosine concentration in unirradiated tumors. Abundance (Mean ± SEM) of (B) myeloid cells (CD45^+^CD11b^+^) and (C) T lymphocytes (CD45^+^TCRb^+^) in 4T1 tumors on days 3,10 after irradiation (8Gy x 3) compared to unirradiated controls.

(A) (B) 4T1-CD45+CD11b+	(C) 4T1-CD45+TCRβ+

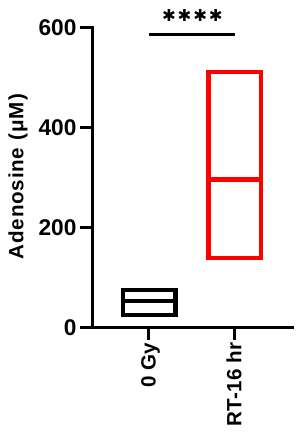

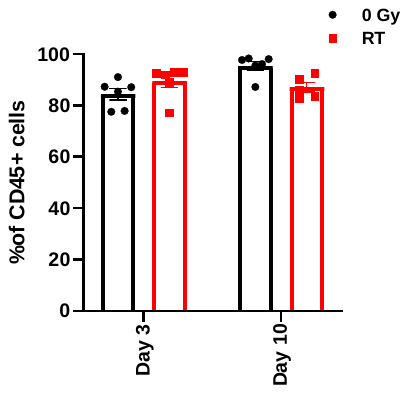

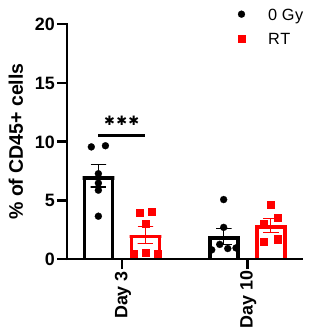

Figure S3: (A) Spider plots of n=6 radiation-treated (8Gy x 3) tumors from LysM^Cre^ mice implanted with EO771 cells (B) Spider plots of n=8 radiation-treated (8Gy x 3) tumors from LysM^Cre^A_2a_R^fl/fl^ mice implanted with EO771 cells. Data are representative of two separate experiments n=10-12 mice/group. (C) slope during tumor growth of radiation-treated (8Gy x 3) tumors from LysM^Cre^ and LysM^Cre^A_2a_R^fl/fl^ mice. (D) Slope (Mean ± SEM) of n=6 radiation-treated (8Gy x 3) tumors from LysM^Cre^ mice implanted with EO771 cells (E) Slope (Mean ± SEM) of n=8 radiation-treated (8Gy x 3) tumors from LysM^Cre^A_2a_R^fl/fl^ mice implanted with EO771 cells.

**(A)	(B)**
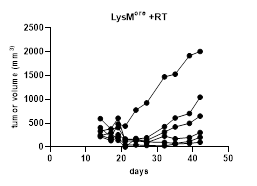

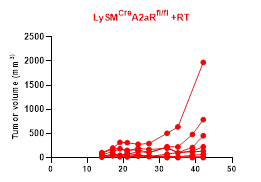

**(C)**
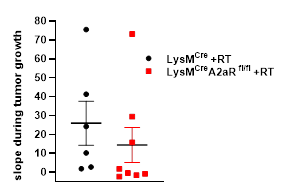

**(D) LysM^cre^ +RT**

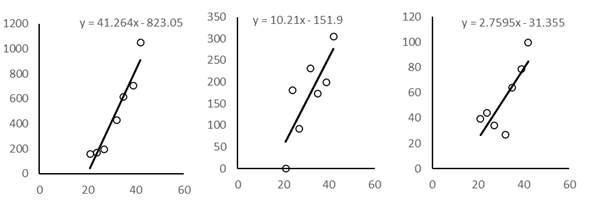

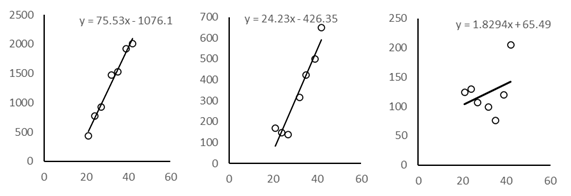

**(E) LysM^Cre^A_2a_R^fl/fl^ +RT**

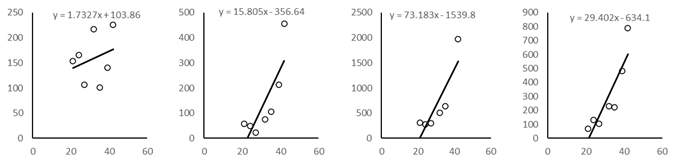

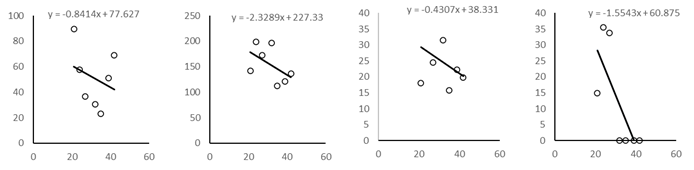

Figure S4: Abundance (Mean ± SEM) of dendritic cells (CD45^+^CD11c^+^), NK cells (CD45^+^CD11b^-^CD11c^-^NK1.1^+^) and B cells (CD45^+^CD11b^-^CD19^+^) in EO771 tumors from mice belonging to all treatment groups on day 11 post-irradiation.

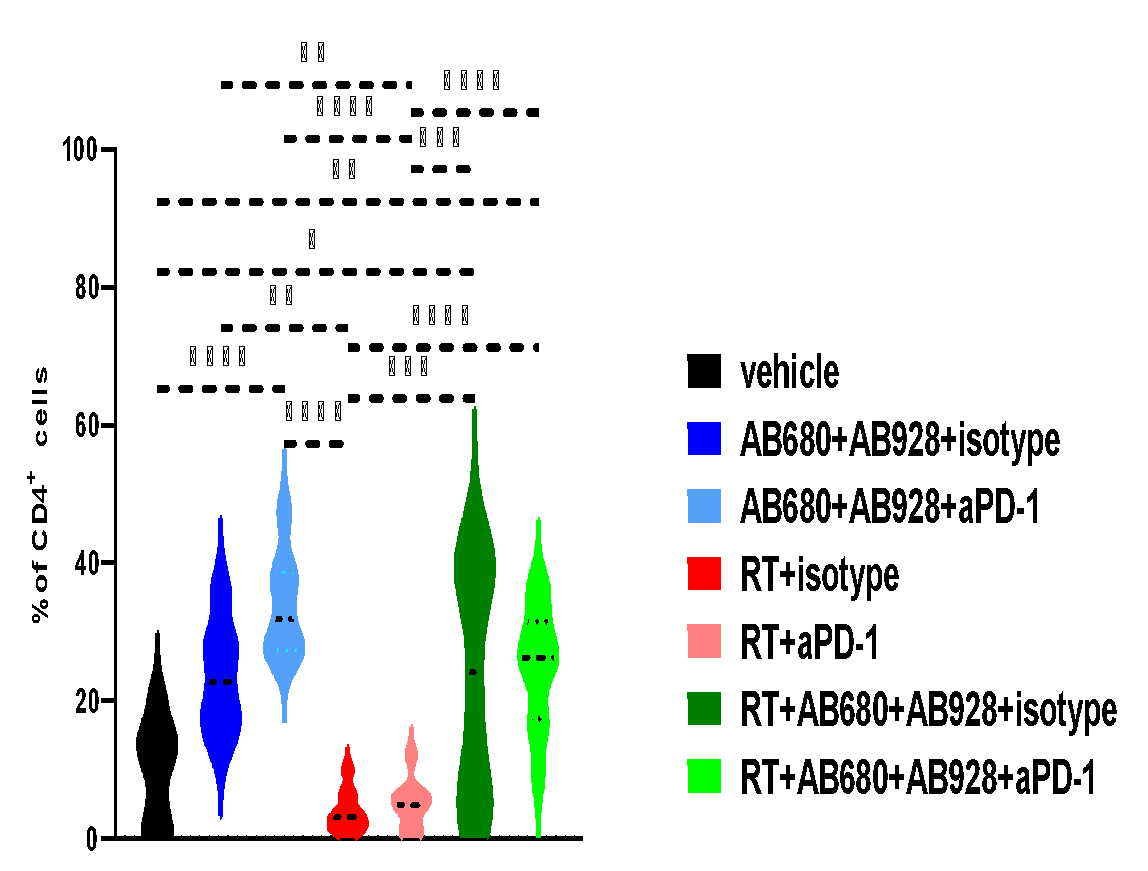

Dendritic cells

NK cells

B cells

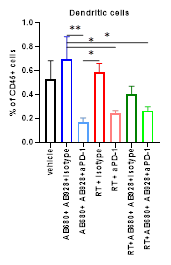

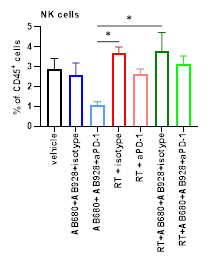

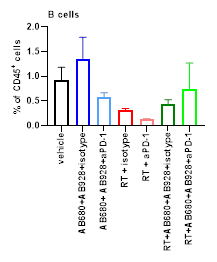

Figure S5: (A) Copy number variation pattern in selected responders and non-responders at pre-therapy, post anti-PD-1 therapy and post anti-PD-1 and SBRT. (B) Abundance (Mean ± SEM) of myeloid cells from responders and non-responders at different stages of therapy.

**(A)**

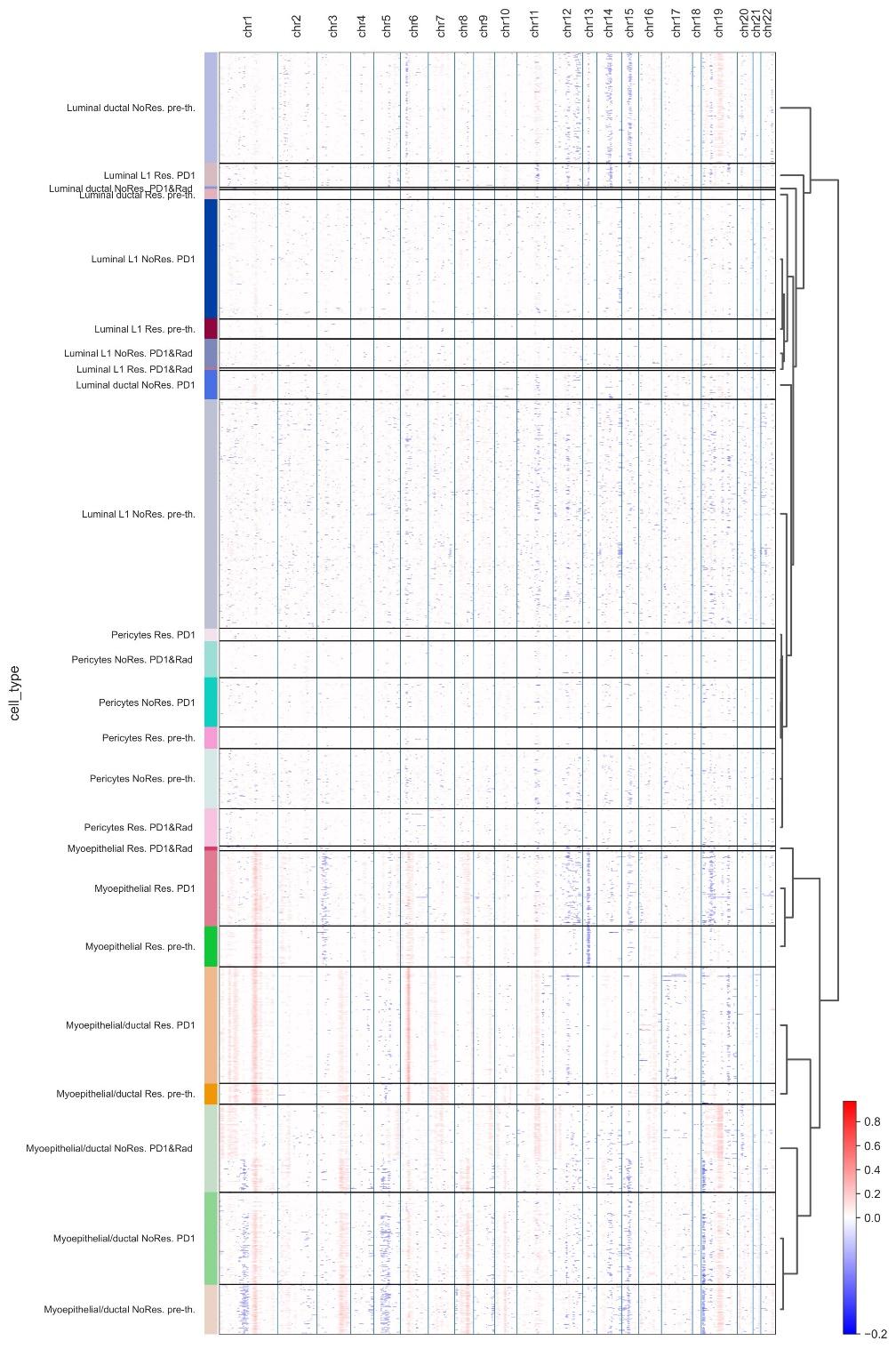

Tumor cells

**(B)**

Responders

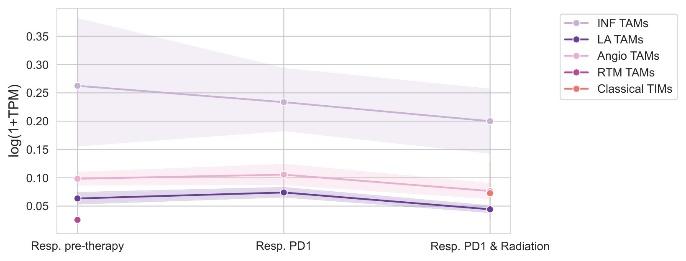

Non-Responders

|  | 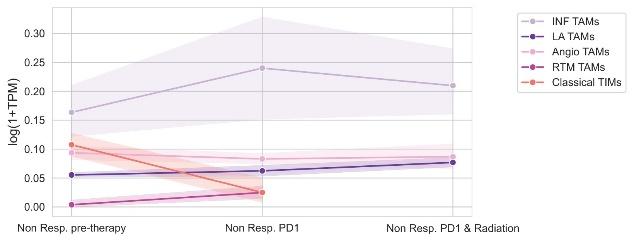 |
| --- | --- |

Supplementary Table 1: Clinical characteristics of breast tumor and adjacent normal tissue collected for adenosine quantification and IHC

| **Patient Number** | **Tumor characteristics** | **Tumor size** | **Procedure** |
| --- | --- | --- | --- |
| 1 | ER+PR+HER2-, Ki67 30% | 9.5 cm X 2.5 cm | Biopsy |
| 2 | ER10%PR>1%HER2:3+, Ki67 40-60% | 3 cm x 6 cm | Biopsy |
| 3 | ER-PR-HER2-, Ki67 35-40% | 2.1cm x 2 cm x2.3 cm | Biopsy |
| 4 | ER+PR-HER2:3+, Ki67 50% | 4 cm | Biopsy |
| 5 | ER-PR-HER2-, Ki67 42% | 2.2 cm | Mastectomy |
| 6 | ER-PR-Her2-, Ki67 75% | 2.1cm | Biopsy |
| 7 | ER95%PR2%HER2-, Ki67 5-10% | 1.2 cm | Partial mastectomy |
| 8 | ER+PR+Her2-, Ki67 25% | 1.1 cm | Mastectomy |
| 9 | ER+PR+Her2-, Ki67 10% | 1.2 cm | Mastectomy |
| 10 | ER+PR+HER2:2+, Ki67 15-20% | 1.1 cm x 0.8 cm x 0.9 cm | Segmentectomy |

Supplementary Table 2: Spectral flow cytometry panel used for immunophenotyping of CD45+ cells in EO771 murine tumors.

| **PHENOTYPING PANEL** | |
| --- | --- |
| **Fluorophore** | **Target** |
| BV421 | enpp1 |
| SB436 | NK1.1 |
| Pac-Blue | Ly6g |
| BV480 | CD103 |
| BV570 | CD19 |
| BV605 | CD11c |
| BV650 | CD4 |
| BV750 | TCRb |
| BV785 | F4/80 |
| A532 | CD45 |
| PE | A_2a_R |
| PE-dazzle594 | CD169 |
| PE-Fire640 | CD8a |
| PE-Cy5 | Ki67 |
| PerCP-Cy5.5 | MHCII |
| PerCPef710 | CD73 |
| PE-Cy7 | FoxP3 |
| APC | Arg1 |
| AF647 | CD39 |
| A700 | CD11b |
| Zombie Near IR | Live/Dead |
| APC-Cy7 | Ly6c |
| APC-Fire810 | B220 |

Supplementary Table 3: Spectral flow cytometry panel used for evaluating T cell effector function in EO771 murine tumors

| **STIMULATORY PANEL** | |
| --- | --- |
| **Fluorophore** | **Target** |
| BV421 | IFN-γ |
| Pac Blue | Granzyme b |
| BV650 | CD4 |
| BV750 | TCRβ |
| A488 | FoxP3 |
| A532 | CD45 |
| PE-Fire640 | CD8a |
| PE-Cy7 | IL-2 |
| APC | TNFα |
| A700 | CD11b |
| Zombie NIR | Live/Dead |

Supplementary Table 4: Differentially expressed genes in CD45^-^, CD45^+^CD11b^+^ and CD45^+^TCRβ^+^ cells and adenosine signaling genes CD45^-^, CD45^+^CD11b^+^ and CD45^+^TCRβ^+^ cells from non-irradiated and irradiated tumors at Day 1, Day 3, Day 5.

| **Cell Type** | **Markers** |
| --- | --- |
| CD45^-^ | COL3A1, TGFB2, CD34, GPA33, FBN1, ECT2, S100A8, S100A9, KIF2C, PTHLH, TGFB1, LUM, ESAM, MMP3, TOP2A, PECAM1, KRT8, KRT18, EMP2, EPCAM, MSLN, ACTA2 |
| CD45^+^CD11b^+^ | PTGS2, MRC1, MERTK, IL1B, SIRPA, FABP5, P2RY14, C1QC, C1QA, CSF3R, SPP1, CXCL9, GPNMB, PPARG, CD33, APOE, ITGAM, ITGAX, CD81, ARG1, CCR2, CCR7, NOS2, CD83, VCAN, LY6C2, LY6G, LY6C1, CD200, CD86, TREM2, H2-AB1, H2-EB1, ADGRE1, CD74, CSF1R, CD14 |
| CD45^+^TCRβ^+^ | CTLA4, PDCD1, SELL, IL2RA, CD44, FOXP3, IL2, TOX, CD8A, CD4, CD69, KLRB1C, NKG7, PRF1, GZMB, TCF7, CD7, GZMA |
| Adenosine signaling | SNAP23, EPB41, CD38, TGFB1, ACTN4, CYTH2, ENPP1, ADORA2A, HP, NECAB2, TSNAX, ADK, NT5E, USP4, ADORA2B, VAMP2, IL10RB, ENTPD1 |

Supplementary Table 5: Patient characteristics of invasive ductal TNBC responders and non-responders selected for the study.

**Responders**

| **Paper**  **Paper number** | **Surgery (Primary)** | **Surgery (Axillary)** | **Original tumor size (cm)** | **Node positive** | **Initial Staging** | **ypCR** | **Residual breast tumor size** | **Post-NAC staging** | **Age** | **Stage** | **Chemotherapy** |
| --- | --- | --- | --- | --- | --- | --- | --- | --- | --- | --- | --- |
| 16 | Lumpectomy |  | 2 | N | T1cN0 | Y |  | ypT0N0 | 33 | IIA | ddACx3 |
| 56 | Mastectomy | SLNB | 2.7 | Y | cT2N1 | Y |  | ypT0N0 | 53 | IIB | ddACx4, Tx12 |
| 46 | Lumpectomy | SLNB | 3 | N | cT2N0 | Y |  | ypT0N0 | 50 | IIA | ACx4, Tx4 |

**Non-responders**

| **Paper number** | **Surgery**  **(Primary)** | **Surgery (Axillary)** | **Original tumor size (cm)** | **Node positive** | **Initial Staging** | **ypCR** | **Residual breast tumor size** | **Post-NAC staging** | **Age** | **Stage** | **Chemotherapy** |
| --- | --- | --- | --- | --- | --- | --- | --- | --- | --- | --- | --- |
| 52 | Mastectomy | SLNB | 4.6 | N | cT2N0 | N | 1.3 | ypT1cN0 | 38 | IIA | TCx12, ACx4 |
| 53 | Mastectomy | ALND | 3.9 | Y | cT2N1 | N | 2.5 | ypT2N1a | 48 | IIB | ACx4, TCx12 |
| 42 | Lumpectomy | SLNB | 2.8 | Y | cT2N1 | N | <0.1 | ypT1aN0 | 79 | IIB | TCx12, ACx4 |

Supplementary Table 6: Markers for identification of CD45^-^ (non-immune), fibroblast, endothelial and CD45^+^ (immune) sub-clusters.

| **Cell Type** | **Markers** |
| --- | --- |
| Luminal L1 | AGR2, AFF3 |
| Myoepithelial | TAGLN, FHOD3 |
| Myoepithelial/ductal | TAGLN, MMP7 |
| Luminal ductal | MMP7, LTF |
| Pericytes | ACTA2, NOTCH3 |
| Fibroblasts | LUM, DCN |
| Endothelial | PECAM1 |
| Inf TAMs | LAMP3 |
| LA TAMs | TREM2, C1QA |
| Angio TAMs | AREG, CCL2 |
| Rtm TAMs | FOLR2, LYVE1 |
| Classical TIMs | S100A4, CLEC11A |
| B cells | MS4A1 |
| Plasmablasts | JCHAIN |
| T cells | CD3D |
| CD8+ T cells | CD8A |
| CD8+ T cells ExRe | CD8A, CXCL13 |
| CD4+ T cells ExRe | CD4, CXCL13 |
| T regs | FOXP3 |
| GmDt T cells | TRDC, XCL1 |

TAMs, Tumor associated macrophages, INF: inflammatory, LA: lipid associated, Angio: angiogenic, Rtm: resident tissue macrophages, ExRe: exhausted reactive, GmD: gamma delta

**Supplementary Methods**

**Maintenance of EO771 and its knockdowns**

EO771 and its knockdowns were maintained in complete Dulbecco′s Modified Eagle′s Medium (DMEM) containing 10% fetal bovine serum (Gibco), 100 U/mL penicillin, 100 mg/ml streptomycin, 2 mM L-glutamine, 1% non-essential amino acids, 1% sodium pyruvate, 1.5 g/l sodium bicarbonate. 4T1 cells were maintained in complete Rosewell Park Memorial Institute media (RPMI) containing 10% fetal bovine serum (Gibco), 100 U/mL penicillin, 100 mg/ml streptomycin, 2 mM L-glutamine, 1% non-essential amino acids, 1% sodium pyruvate.

**Tumor harvest**

On the day of harvest, mice were euthanized, tumors were harvested, weighed, and finely diced in 6 well plates containing serum free RPMI containing DNase (40 μL of 20 mg/mL solution; Roche; Basel, Switzerland) and collagenase D (125 μL of 40 mg/mL solution; Roche; Basel, Switzerland). Tumor samples were then incubated on a shaker at 37°C for 30min. Tumor digests were quenched with 5ml complete RPMI, physically mashed and filtered through 70μm filters (Miltenyi; Bergisch Gladbach, GE) and pelleted. RBCs were lysed by incubating digests at room temperature for 4 min with 4 ml ACK buffer (Quality Biological; Gaithersburg, Maryland, USA) and then quenched by topping off with PBS prior to pelleting. All samples were then placed in U-bottom 96-well plates for staining.

**Immunohistochemistry**

Formalin-fixed, paraffin-embedded (FFPE) sections were generated at an optimal thickness of 3-4 µm. The sections were then deparaffinized and rehydrated prior to antigen retrieval using 0.01 M citrate buffer (pH 6). Slides were then washed, followed by quenching of endogenous peroxidase with 3% hydrogen peroxide for 10 min and then washed. Thereafter, sections were blocked with 10% horse serum for 30 min and then incubated with primary antibodies against A_2a_R (clone 7F6-G5-A2, Abcam, Cat#: ab79714, RRID: AB_1603112) at 1:4000 or against CD73 (clone D7F9A, Cell signaling technologies, Cat#: 13160S, RRID: AB_3750601) at 1:100 for 1.5hr at RT. After washing, secondary antibody (Horse anti-mouse biotinylated, Vector# BA2000) was added, and slides were incubated for 30 min at room temperature. After washing, avidin-biotin complex (Vector #PK6100) was added to the slides for 30 min. Thereafter, DAB solution was added to each section and subsequently, slides were counterstained with hematoxylin and dehydrated in ethanol followed by mounting for reading under a light microscope. The expression of CD73 and A_2a_R in cancer cells and immune cells was reported using the H-scoring system. The H-score is calculated by combining the percentage of cells with a specific staining intensity (0=no staining, 1=weak, 2=moderate, 3=strong) with a weighted factor for each intensity level.

**Quantification of adenosine signaling genes**

RT-qPCR was used to determine the amount of adenosine signaling genes in EO771 and 4T1 cells. RNA was extracted from EO771 and 4T1 cells using RNeasy Plus Mini Kit (Qiagen) following the manufacturer's protocol. cDNA synthesis was generated using Superscript IV first strand synthesis kit (ThermoFisher Scientific) followed by amplification using TaqMan Universal Master Mix II (Applied Biosystems). Commercially available TaqMan primers from ThermoFisher Scientific were used for CD73 (Mm00501910_m1), CD39 (Mm00515447_m1), ENPP1 (Mm00501097_m1), A_2a_R (Mm00802075_m1), A_2b_R (Mm00839292_m1). Reactions were run on the Quant-Studio 6 platform (Applied Biosystems) with an initial denaturation step at 95°C for 10 min and 40 cycles of denaturation at 95°C for 15 sec followed by annealing/extension at 60°C for 1 min. Relative quantification of adenosine signaling genes was determined by calculating 2^-ΔCt^ with respect to the housekeeping gene, GAPDH.

**Flow cytometry**

Samples were washed with PBS. All staining and fixation steps were performed for 30 min at room temperature protected from light. Dead cells were stained by resuspension in 100μL PBS+Live/Dead Fixable Blue dye (1:500; Invitrogen; Waltham, Massachusetts, USA). Samples were washed twice PBS and resuspended in FACS (PBS+3%FBS+1mM EDTA+10mM HEPES) supplemented with TruStain FcX (1:50; BioLegend; San Diego, California, USA) and placed on ice. Surface antibodies (supplementary table 2) were prepared at optimal dilutions in FACS supplemented with Brilliant Stain Plus buffer (BD; Franklin Lakes, New Jersey, USA) and added to samples in blocking solution. Post staining, samples were washed twice with FACS buffer and fixed in 100ul of FoxP3 Fixation/Permeabilization Kit buffer (eBioscience; San Diego, California, USA). Samples were washed twice with 1X Permeabilization buffer (1X PW) and then stained in 1X PW plus intracellular antibodies (supplementary table 2). Samples were then washed twice in 1X PW then fixed in 100μL of FluoroFix buffer (BioLegend), washed twice with FACS, then resuspended in 200 μL FACS and acquired on a Cytek Aurora 3-laser cytometer. Simultaneously stained splenocyte samples were used for single stain controls. For studying T cell effector function, *ex vivo* tumor infiltrating lymphocyte stimulation was performed followed by intracellular cytokine staining. Briefly, single cell suspensions were made from tumors as described above. 5-10 x 10^6^ cells were resuspended in 50ul complete RPMI and plated in 96 well round bottom plate with 50ul complete RPMI containing phorbol myristate acetate (PMA) and ionomycin (InvivoGen) at a final concentration of 50 ng/ml and 500 ng/ml. After 30 min, 100ul of complete RPMI containing PMA/ionomycin and Brefeldin A at a final concentration of 5 μg/mL (BioLegend) were added. After 3.5 hours (4 hours total stimulation), cells were washed and fixed with the FoxP3 Fix/Perm Kit (eBioscience) for 30 to 60 minutes on ice in the dark. Cells were then stained for surface and intracellular markers described in supplementary table 3 and acquired on a Cytek Aurora 3-laser cytometer.

**Bulk RNA sequencing**

Tumors were harvested on day 1, day 3 and day 5 post RT (8 Gy X 3), dissociated, and single cell suspensions were made. Cells were stained with PI (ThermoFisher Scientific), DRAQ5 (ThermoFisher Scientific), BUV395 anti-CD45 (BioLegend), FITC-CD11b (BioLegend), BV750-TCRβ (BioLegend). CD45^-^, CD45^+^CD11b^+^, CD45^+^TCRβ^+^ cell populations were sorted on a FACS Aria (BD Biosciences) followed by RNA isolation using Qiagen RNAeasy Mini Kit (Qiagen). Paired end bulk RNA sequencing (RNA-seq) was performed on sorted immune cell subsets by Azenta Life Sciences, USA. Sequence quality was assessed using fastQC (RRID:SCR_014583) and STAR (RRID:SCR_004463) was used to index genome and BAM generation. Sequence reads were aligned to the mouse reference genome GRCm39 from Ensembl (RRID:SCR_002344) (31), and the count matrix was generated by featureCounts (RRID:SCR_012919). Above-mentioned RNA sequence processing steps were conducted on Columbia University C2B2 HPC cluster. Gene expression counts of genes listed in supplementary table 4 were normalized to counts per million (CPM). For each cell compartment and time point, log2 fold changes were calculated by subtracting log2(CPM+1) expression in the untreated sample from the corresponding irradiated sample and visualized in R (<http://www.rstudio.com/>, RRID:SCR_000432).

**Single cell RNA sequencing**

From the phase Ib/II trial offering women with operable triple negative breast cancer (TNBC) neoadjvuant pembrolizumab and whole breast irradiation followed by chemotherapy (ddAC, T) (NCT03366844, 26), we identified three patients who achieved complete pathologic responses (responders) and three that did not (non-responders) at the time of lumpectomy or mastectomy. Inclusion criteria for patients included- age matched responders and non-responders bearing non-metastatic tumors > 2cm, T2+N0-1M0 who had high quality scRNA sequencing data from both CD45^-^ and CD45^+^ cells deposited by Shiao et al (26). Patients received pembrolizumab at a dose of 200 mg, administered intravenously, on day 1 and 21 and received three daily fractions of 8 Gy stereotactic radiation to the breast tumor between day 21 and 28. Biopsies were obtained at (1) baseline, pre-treatment, (2) 3 weeks after starting pembrolizumab, and (3) 3 weeks after completing radiotherapy with concurrent pembrolizumab. Neoadjuvant chemotherapy was initiated by day 35 (after all biopsies were obtained) for all patients. After completion of chemotherapy, patients proceeded to lumpectomy or mastectomy (26). Additional details regarding trial design and characteristics of patients used for our research are available in supplementary table 5. The scRNA sequencing data analyzed from selected patients in this study were obtained from Gene Expression Omnibus (GEO, RRID:SCR_005012) at GSE246613.

**ScRNA-seq analyses**

scRNA sequencing (scRNAseq) data generated from CD45- (non-immune) and CD45+ (immune cells) compartment from 3 responders (paper number 16, 56, 46; supplementary table 5) and 3 non-responders (paper number 52, 53, 42; supplementary table 5) was analyzed. Cell-gene matrices were pre-processed by excluding cells with fewer than 500 detected genes. Cells with more than 10% of transcripts originating from mitochondrially encoded genes were also removed. Expression matrices were then normalized using log₂ (1+TPM), where TPM represents transcripts per million. Latent space representations of the datasets were inferred using an approach based on Random Matrix Theory (RMT) (32). Clustering was performed using the Leiden algorithm, as implemented in (33, 34), leveraging the latent space derived from the RMT framework. The optimal number of clusters was determined by maximizing the mean silhouette score across a range of Leiden resolutions. Specifically, multiple clustering iterations were conducted, and the resolution yielding the highest mean silhouette score was selected. In some cases, sub-clustering of specific populations was performed by reapplying this procedure within a given cluster, which proved particularly useful for resolving immune populations. The robustness of sub-clustering was validated through supervised visualization of known marker genes associated with these populations. Genes highlighted in the dot plots were selected based on differential expression analysis. Candidate genes were required to pass a t-test (target population versus all others) with a Benjamini–Hochberg corrected p-value < 0.01. As an additional criterion, selected genes had to be expressed in at least 60% of cells within the target population and in fewer than 25% of cells across all other populations. Gene signatures for each human epithelial population were similarly generated, selecting all genes with a corrected p-value ≤ 0.05, while maintaining the same expression thresholds of ≥ 60% within the target population and ≤ 25% across others.

To visualize single-cell clusters, dimensionality reduction was performed using uniform manifold approximation and projection (UMAP). Default parameters were applied for UMAP, 15 neighbors and a minimum distance of 0.3. Visualizations, and SCANPY (32). InferCNV (34) was applied to the scRNA-seq datasets to identify malignant epithelial cells exhibiting genomic instability. Epithelial cells classified as non-malignant based on inferred copy number alterations (CNVs) may represent either true benign cells or transformed cells lacking detectable CNVs by scRNA-seq. Each patient sample was first analyzed independently using transcriptomic denoising and clustering approaches, as described above, to identify populations corresponding to those in the human consensus atlas. InferCNV was then run within each cohort to detect cell populations harboring CNVs.

**Single cell dimensionality reduction and clustering of non-immune and immune libraries**

In the CD45^-^ compartment, clusters were annotated as luminal L1, myoepithelial, myoepithelial/ductal, luminal ductal, pericytes based on markers in supplementary table 6. Similarly, in the CD45+ compartment, clusters were annotated as inf TAMs, LA TAMs, angio TAMs, rtm TAMs, classical TIMs, B cells, plasmablasts, T cells, CD8+ T cells, CD8+ T cells ExRe, CD4+ T cells ExRe, Tregs, gamma delta T cells based on markers in supplementary table 6. Endothelial and fibroblast sub-clusters were annotated based on markers in supplementary table 6. Each sample and patient dataset was analyzed independently and subsequently integrated using population-specific genes identified through differential expression analysis, as described above. We utilized the cell populations identified through the analysis described in the previous section to generate line plots, which reflect the mean expression values (represented by dots in the figures) and also the 95% confidence intervals (bands in the figures) corresponding to the true distribution of all cells within each population.
